# Organization, development, and plasticity of the serotonergic dorsal raphe nucleus projectome

**DOI:** 10.64898/2026.03.04.709518

**Authors:** Lucía Jiménez-Fernández, Zoltán Kristóf Varga, Klara Thurner, Florence Kermen

## Abstract

The dorsal raphe nucleus (DRN) is the main source of serotonin (5-HT) in the central nervous system. 5-HT DRN neurons modulate a wide range of physiological functions and brain states via their widespread and heterogeneous projections. Yet the organizational logic, developmental dynamics, and experience-dependent plasticity of these projections remain poorly understood.

By reconstructing the whole brain projections of 81 individual 5-HT DRN neurons across developmental stages in the larval zebrafish, we find that these neurons exhibit diverse projection patterns, ranging from broadly distributed to spatially confined. We identify two previously uncharacterized, projection-defined subpopulations within the larval dorsal raphe nucleus, which selectively target either the telencephalon or the spinal cord. This organization is consistent with the existence of anatomically distinct serotonergic pathways that may differentially influence forebrain and sensorimotor circuits. Longitudinal analysis revealed that the gross architecture of 5-HT neurons remained stable during larval development, although region-specific remodeling occurred in all neurons. Importantly, stress exposure during the first week of development, but not thereafter, altered the density of 5-HT DRN projections to telencephalic regions, suggesting a critical window of increased plasticity during early life.

Collectively, our findings clarify the developmental trajectory, maturation, and stress-induced plasticity of the serotonergic circuitry, providing important insights for deciphering the pathophysiology of stress-related mental disorders.

## Introduction

The dorsal raphe nucleus (DRN) is the main source of serotonin (5-HT) in the central nervous system (CNS) ^1,2^. Through its extensive projections, the DRN modulates a wide range of physiological functions and brain states, including locomotion, arousal, stress responses, social interactions, and anxiety-like behaviors ^3–7^. The functional diversity of the DRN arises partly from the remarkable heterogeneous nature of 5-HT neurons, which differ in morphology, gene expression, neurotransmitter phenotype, and connectivity ^8–11^. Characterizing how this neuronal diversity is organized within the DRN and how it emerges during early development is key to understanding the mechanisms through which the serotonergic system shapes brain function and behavior in early life.

Serotonergic neurons are among the earliest developing neuromodulatory populations in the vertebrate brain. The first neurites of 5-HT DRN neurons emerge during embryonic stages, shortly after differentiation ^12,13^, and by early developmental stages, they already innervate large portions of the CNS. This early innervation coincides with critical periods of circuit assembly, synaptogenesis, and behavioral emergence. Serotonin plays a key developmental role during this phase, modulating axon guidance, dendritic growth, and synaptic maturation ^14,15^. While the timeline of developmental serotonergic innervation has been broadly characterized ^12,13,16^, how individual 5-HT DRN neurons establish and refine their projections during early life remains unexplored.

Early life is also a period of heightened sensitivity to environmental influences, and experiences during this stage can have lasting effects on brain function and behavior. Specifically, stress exposure during critical developmental windows can disrupt the maturation of serotonergic circuits and their targets, and impair brain-wide functional connectivity in the DRN ^17–20^. However, it remains unclear whether early life stress (ELS) exposure reshapes the anatomical organization of the 5-HT DRN projectome and whether such effects are restricted to specific developmental windows of plasticity.

Here, we map the 5-HT DRN projectome in zebrafish larvae at single-neuron resolution to dissect its organization, development, and susceptibility to ELS. We identify distinct subpopulations of 5-HT neurons based on their projection profiles, targets, and organization within the DRN. We find that the organization of the 5-HT DRN projectome remains largely stable during early development, although individual neurons undergo notable neurite remodeling, with some brain regions exhibiting more pronounced structural changes. Finally, we show that ELS induces region-specific alterations in DRN projections during a temporally restricted window of development. Together, these findings provide a developmental framework for understanding the organization and plasticity of the serotonergic DRN, revealing how a major neuromodulatory system emerges, stabilizes, and adapts during early life.

## Results

### 5-HT DRN neurons provide highly heterogeneous projections across brain regions

We established a whole-brain map of the 5-HT DRN projectome at single-cell resolution in one-week-old larvae. We selected sparsely labeled Tg(*tph2:gal4:UAS:caax-GFP;gadb1:DsRed*) fish with GFP expressed at the membrane of a single neuron within the DRN, performed whole-brain confocal imaging, traced neurites and registered them onto a template brain (Fig. 1A). The projections of 81 5-HT DRN neurons were reconstructed (Sup. Fig. 1), which exhibit variable projection profiles (Fig. 1B) and collectively innervate a large part of the central nervous system (CNS) (Fig. 1C), similar to the complete DRN projection pattern (Sup. Fig. 2 A-D).

**Figure 1:**
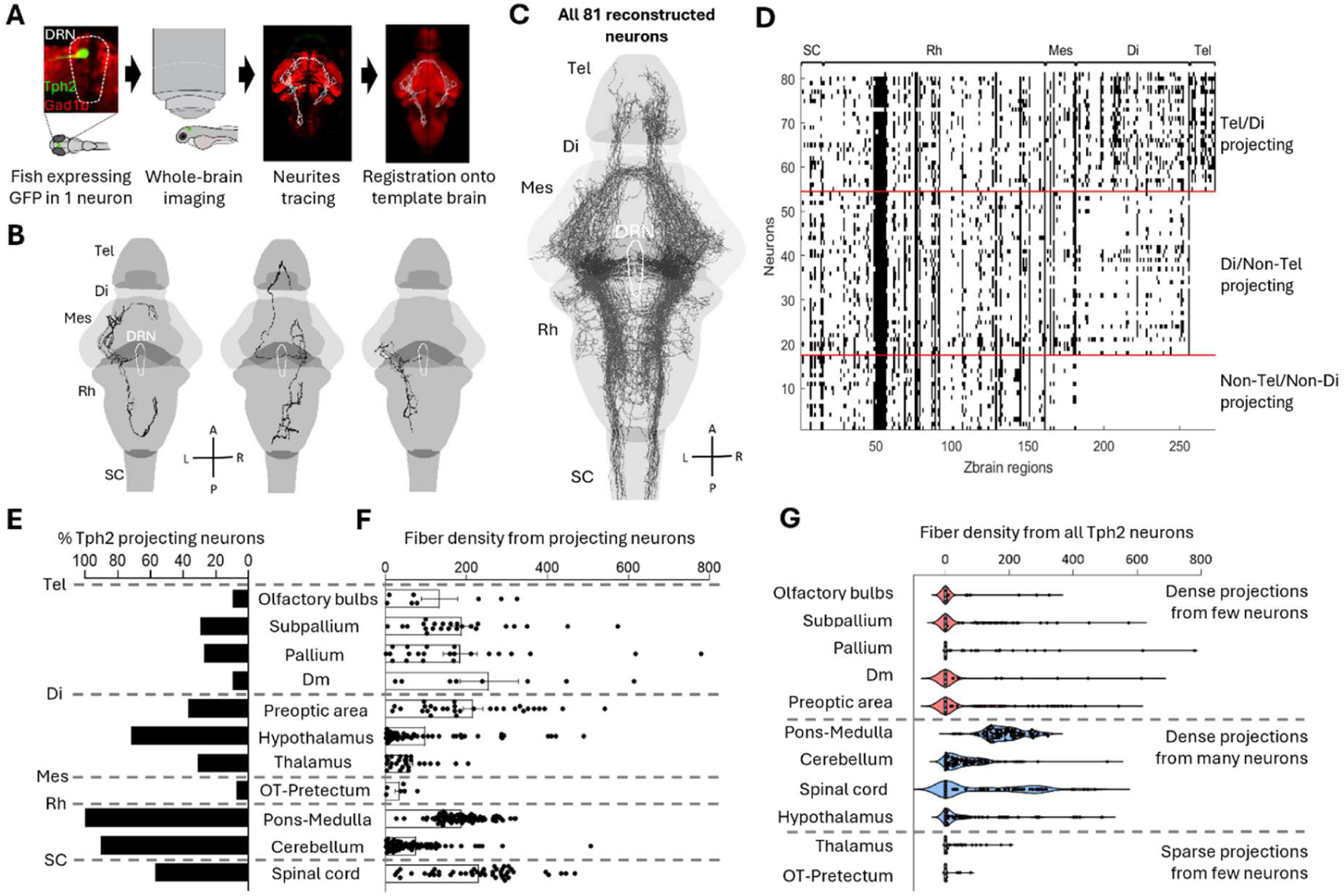
Mapping and classification of 5-HT DRN fiber density across brain regions. **A.** Schematic representation of the single-neuron tracing pipeline. **B.** Representative examples of individual Tph2 DRN neurons reconstructions with diverse projection patterns. **C.** Overlay of all 81 reconstructed Tph2 DRN neurons. **D**. Matrix of innervated Zbrain regions by all neurons. Black=fibers, white=no fibers. **E.** Percentage of Tph2 DRN neurons projecting to selected regions of the central nervous system. **F.** Density of Tph2 DRN neurons projections to selected regions of the central nervous system (mean ±SEM). Density = 4.10^-9^ fiber’s area/ brain region area. **G.** Distribution of projection density for all reconstructed Tph2 neurons, distinguishing the three categories of regions identified by the zero-inflated model. Density = 4.10^-9^ fiber’s area/ brain region area. DRN: Dorsal raphe nucleus, Tel: Telencephalon, Di: Diencephalon, Mes: Mesencephalon, Rh: Rhombencephalon, SC: Spinal cord, OT: Optic tectum, Dm: Dorsomedial telencephalon A: Anterior, P: Posterior, L: Left, R: Right.

**Figure 2:**
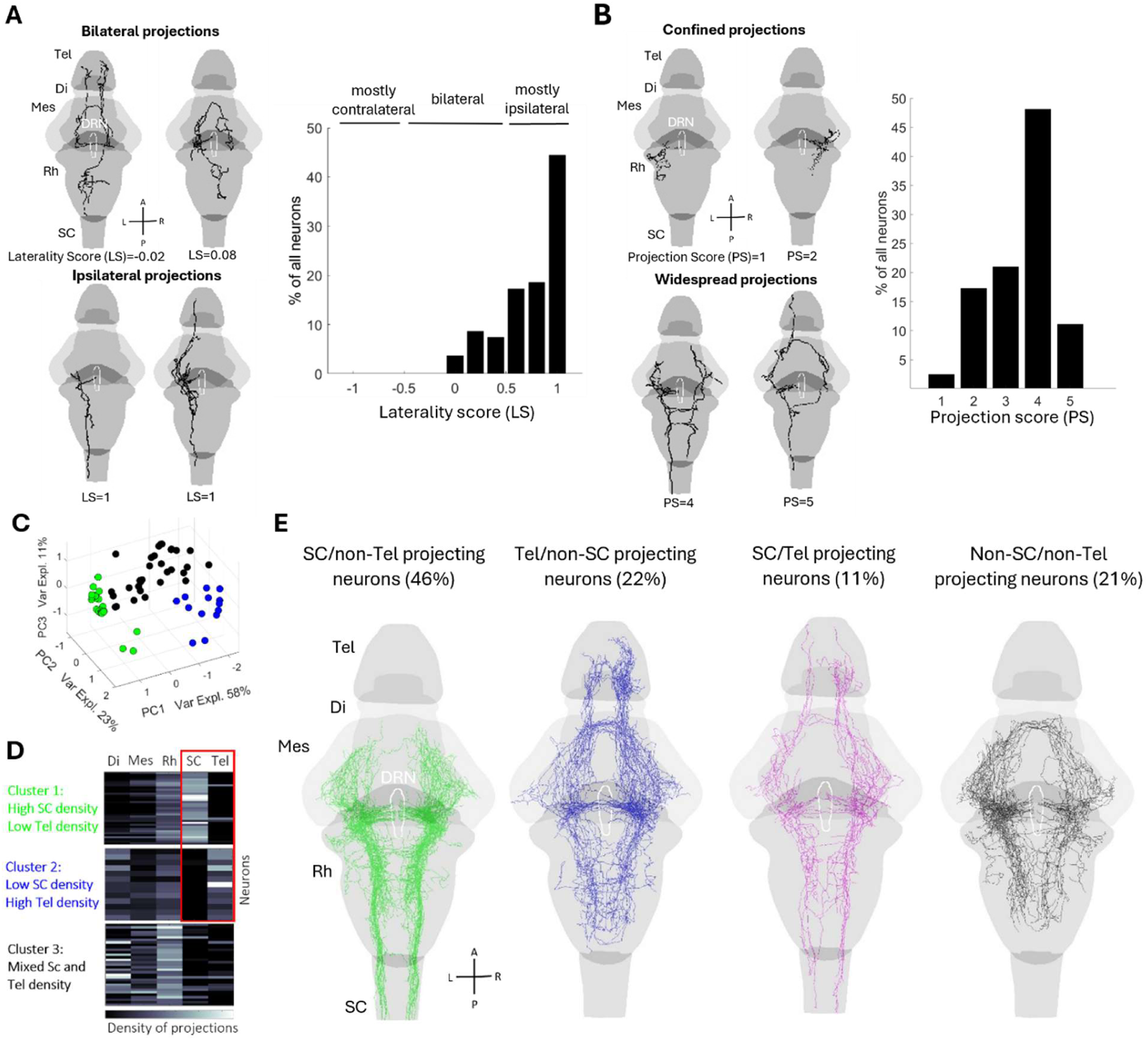
Heterogeneity of the 5-HT DRN projection patterns. **A.** Examples of Tph2 neurons that project bilaterally (left, top) and exclusively to ipsilateral regions (left, bottom), and laterality score of all Tph2 neurons (right). Laterality score = (ipsilateral − contralateral)/(ipsilateral + contralateral); laterality value of 1 indicates purely ipsilateral; 0 indicates equal amount of contralateral and ipsilateral projections and −1 indicates purely contralateral projections. **B.** Examples of Tph2 neurons that display confined projections (left, top) and widespread projections (left, bottom), and projection score of all Tph2 neurons. Projection score indicates the number of main CNS subdivisions innervated. **C.** Projection density to the five brain subdivisions onto the first three PCs. **D**. Projection density to the five brain subdivisions by the three clusters. Black=no fibers, white=high fiber density. **E.** Overlay of all Tph2 neurons projecting to spinal cord but not to telencephalon (green; SC/non-Tel), to telencephalon but not to spinal cord (blue; Tel/non-SC), to both (magenta; Tel/SC), or to neither (black; non-SC/non-tel). DRN: Dorsal raphe nucleus, Tel: Telencephalon, Di: Diencephalon, Mes: Mesencephalon, Rh: Rhombencephalon, SC: Spinal cord, LS: Laterality score, PS: Projection score A: Anterior, P: Posterior, L: Left, R: Right.

To examine how brain regions differ in terms of the 5-HT DRN projections they receive, we calculated the Tph2^+^ fiber density in 294 previously delineated brain regions in the Zbrain atlas ^21^ (Fig. 1D). Since these include overlapping regions, and peripheral nervous system areas devoid of 5-HT fibers, we focused on 11 CNS regions known to receive 5-HT fibersin zebrafish ^4,13,22–24^, and which cover 80.3% of the CNS depicted in the Zbrain atlas.

Some brain regions, such as the cerebellum and hypothalamus, were targeted by over 70% of the neurons, while others, like the olfactory bulbs and optic tectum, received projections from fewer than 10% (Fig. 1E). To quantify regional differences in serotonergic projections, we calculated the average fiber density for each region, focusing only on neurons that project there (Fig. 1F). The spinal cord and telencephalic brain regions exhibited a high projection density, whereas the thalamus and optic tectum were sparsely innervated. To objectively categorize brain regions based on their innervation by individual 5-HT DRN neurons, we used a zero-inflated model that captures differences arising from both the proportion of projecting neurons and the density of inputs received. This analysis revealed three categories of brain regions (Fig. 1G): telencephalic areas and preoptic area are densely innervated by only a few neurons; pons-medulla, cerebellum, spinal cord, and hypothalamus are densely innervated by many neurons; thalamus and optic tectum are sparsely innervated by few neurons. Together, these results reveal a diverse serotonergic architecture in which different brain regions receive highly variable input from DRN neurons.

### Organizational logic of 5-HT DRN neurons’ heterogeneous projections

The first indication of the diversity of DRN neurons in our data consists of the laterality of their projections, i.e., the extent to which their neurites project to the ipsilateral versus contralateral hemisphere relative to the soma. Some neurons remain exclusively ipsilateral, while others exhibit both contralateral and ipsilateral projections (Fig. 2A, left). Overall, most neurons innervate both hemispheres, although their projections are predominantly ipsilateral (Fig. 2A, right). Only 3% of the neurons display equal amount of contralateral and ipsilateral fibers, and none of the neurons project preferentially to the contralateral side.

Another feature of the diversity of the 5-HT DRN neurons’ projections is their reach, i.e., how widespread or confined their projections are (Fig. 2B, left). More than 60% of the neurons exhibit extensive projections to at least three of the five CNS subdivisions, whereas only 20% of the neurons display spatially confined projections, innervating only one or two subdivisions (Fig. 2B, right).

To assess whether subsets of 5-HT DRN neurons exhibit broad similarity in projection patterns, we performed hierarchical clustering and principal component analysis to group neurons based on their projection density to the five CNS subdivisions. This analysis revealed three distinct neural clusters (Fig. 2C). Cluster 1 showed high spinal cord and low telencephalon fiber densities, while cluster 2 exhibited the opposite pattern (Fig. 2D). To investigate this further, we directly categorized all neurons based on whether they project to the telencephalon or the spinal cord, independent of the PCA-based clustering analysis (Fig 2 E). Only 11% of 5-HT DRN neurons project to both the telencephalon and spinal cord, while 68% innervate only one of these two regions (46% spinal cord-not telencephalon, 22% telencephalon-not spinal cord; Fig. 2E). Therefore, our results reveal two projection-based subsets of 5-HT DRN neurons with largely mutually exclusive projections to either the telencephalon or the spinal cord.

Prior evidence in mice ^25^ and zebrafish ^26^ suggests that raphe neurons can be subdivided according to their rhombomeric origin, highlighting the importance of examining whether the anteroposterior position of neuronal soma influences their projection patterns. To determine whether 5-HT neurons’ projection pattern depends on their soma location within the DRN, we quantified the projections distribution along the anterior–posterior and dorso–ventral axes using anteriority and ventrality indices (Fig. 3 A-D, Supp. table 1, Methods). While no global topographic organization was found based on the soma’s dorso-ventral position, neurons in the anterior DRN project preferentially to ventral regions (Fig. 3 B).

**Figure 3:**
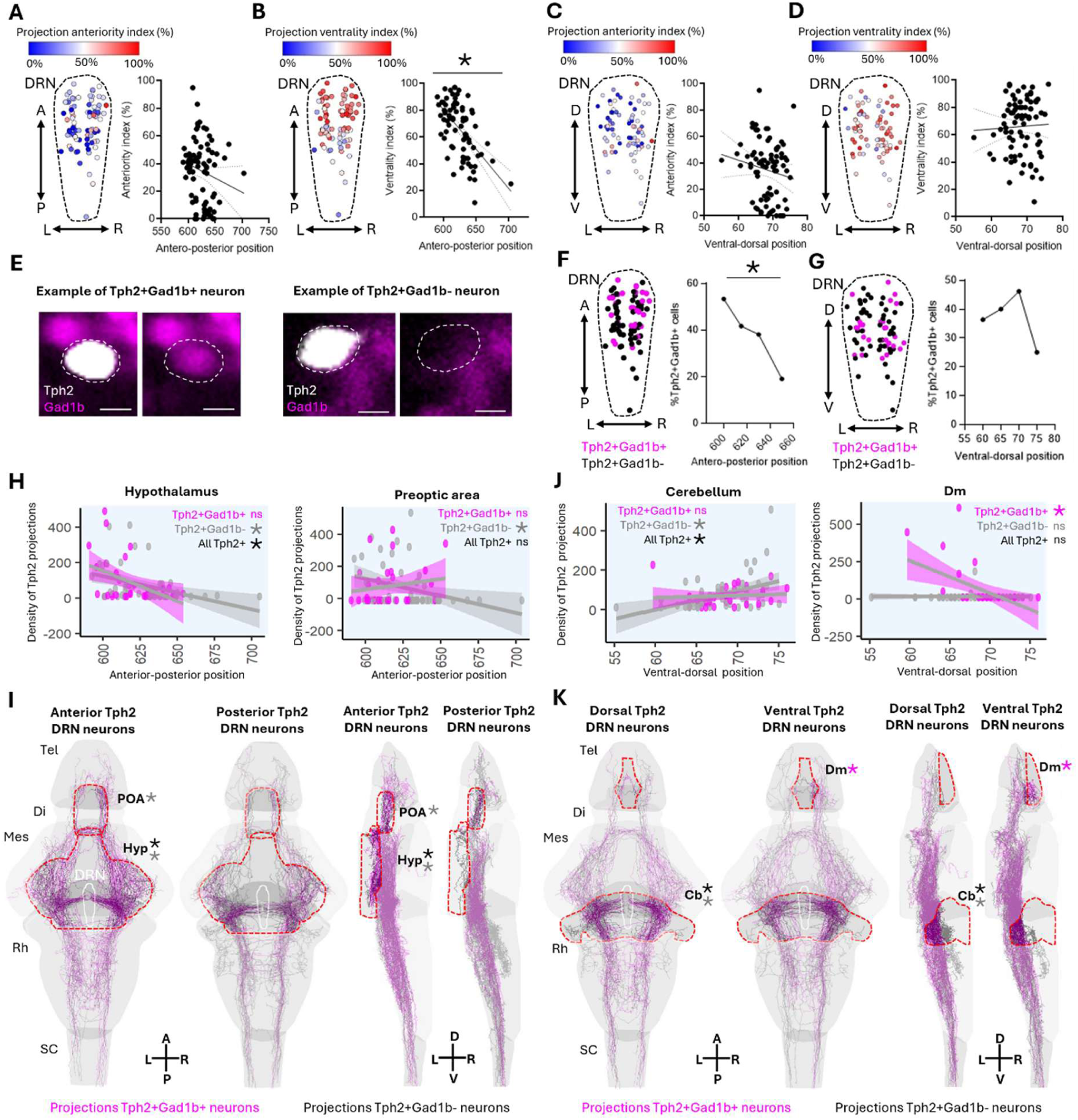
Topographical organization of the 5-HT DRN projectome. **A.** Anteriority index of Tph2 neurons’ projections across the anterior-posterior DRN axis (Spearman correlation: n=81, rho=-0.169, p=0.131). The anteriority index indicates the proportion of a neuron’s neurite that is located anterior to its soma. **B.** Ventrality index of Tph2 neurons’ projections across the anterior-posterior DRN axis (Spearman correlation: n=81, rho=-0.558, p<0.0001). The ventrality index indicates the proportion of a neuron’s neurite that is located ventral to its soma. **C.** Anteriority index of Tph2 neurons’ projections across the dorsal-ventral DRN axis (Spearman correlation: n=81, rho=-0.171, p=0.127). **D.** Ventrality index of Tph2 neurons’ projections across the dorsal-ventral DRN axis (Spearman correlation: n=81, rho=0.041, p=0.712). **E.** Examples of Tph2^+^Gad1b^+^ and Tph2^+^Gad1b^-^ neurons. Scale bar = 5 µm. **F.** Distribution of Tph2^+^Gad1b^+^ and Tph2^+^Gad1b^-^ neurons across the antero-posterior DRN axis (Spearman correlation: n(gad1b^+^, gad1b^-^)=30,51, rho=-0.302, p=0.006). **G.** Distribution of Tph2+/Gad1b+ and Tph2+/Gad1b-neurons across the dorsal-ventral DRN axis (Spearman correlation: n(gad1b^+^, gad1b^-^)=30,51, rho=0.027, p=0.812). **H.** Correlation between anterior-posterior soma position and fiber density in hypothalamus (Spearman correlation all neurons: n=81, rho=-0.427, p=0.002; Spearman correlation gad1b^-^ neurons: n=51, rho=-0.509, p=0.004) and preoptic area (Spearman correlation gad1b-neurons: n=51, rho=-0.407, p=0.033). Density = 4.10^-9^ fiber’s area/ brain region area. **I**. Whole-brain projection map of anterior and posterior Tph2 DRN neurons. **J.** Correlation between ventral-dorsal soma position and fiber density in cerebellum (Spearman correlation all neurons: n=81, rho=0.390, p=0.005; Spearman correlation gad1b^-^ neurons: n=51, rho=0.449, p=0.016) and Dm (Spearman correlation gad1b^+^ neurons: n=30, rho=-0.566, p=0.036). Density = 4.10^-9^ fiber’s area/ brain region area. **K.** Whole-brain projection map of dorsal and ventral Tph2 DRN neurons. DRN: Dorsal raphe nucleus, Tel: Telencephalon, Di: Diencephalon, Mes: Mesencephalon, Rh: Rhombencephalon, SC: Spinal cord, Hyp: Hypothalamus, POA: Preoptic area, Cb: Cerebellum, Dm: Dorsomedial telencephalon, A: Anterior, P: Posterior, L: Left, R: Right.

In line with recent studies^24,27^, we found 36% of Tph2^+^Gad1b^+^ neurons, which are mainly located in the anterior DRN (Fig. 3E-G, Supp. table 1). Region-specific analyses revealed that anterior DRN neurons innervate the hypothalamus and preoptic area more than posterior neurons (Fig. 3H-I, Supp. table 1). Dorsal neurons provide denser projections to the cerebellum, and Tph2⁺Gad1b⁺ neurons, mainly from the ventral DRN, dominate Dm projections (Fig. 3J–K, Supp. table 1).

### Stability and regional remodeling of the developing 5-HT DRN projectome

We tracked developmental changes in the 5-HT DRN projectome by longitudinally imaging Tg(*tph2:gal4:UAS:caax-GFP;gadb1:DsRed*) fish at 1 and 2 weeks and reconstructing 19 individual 5-HT DRN neurons to identify pruned and new neurite segments (Fig. 4A, Sup. Fig. 4).

**Figure 4:**
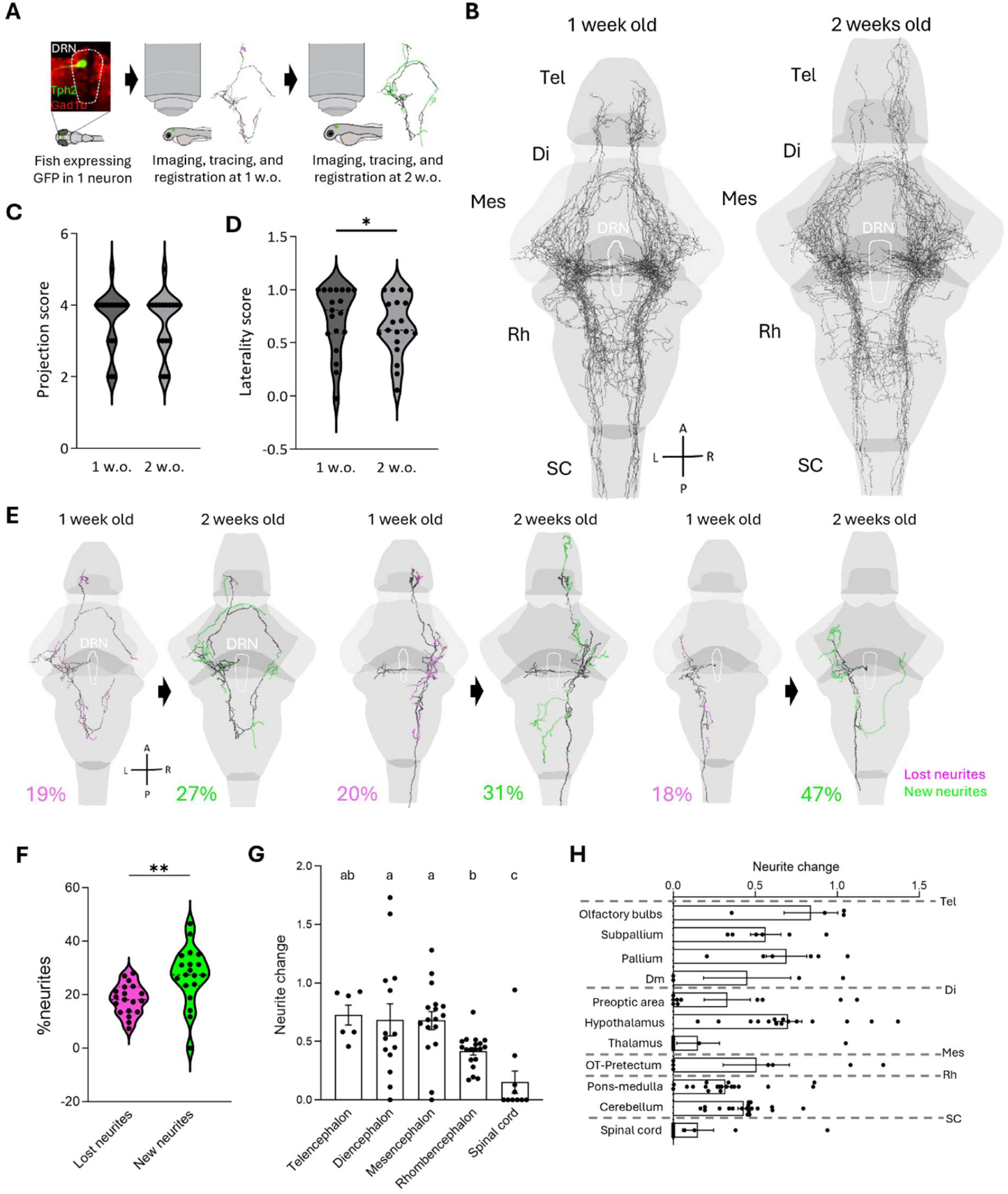
Developmental remodeling of the 5-HT DRN projections in early life. **A.** Schematic representation of the single-neuron tracing pipeline for reconstruction of the same Tph2 DRN neurons at 1 week and 2 weeks of age. **B.** Overlay of all 19 reconstructed Tph2 neurons at 1 week and 2 weeks old. **C.** Projection score at 1 week and at 2 weeks. Projection score indicates the number of CNS subdivisions innervated (paired t-test: n=18, p=0.331). **D.** Laterality score at 1 and 2 weeks old (paired t-test: n=19, p=0.021). Laterality score = (ipsilateral − contralateral)/ (ipsilateral + contralateral); values range from 1 for purely ipsilateral to 0 for fully bilateral projections. **E.** Three examples of neurons reconstructed at 1 week and 2 weeks old. Magenta: pruned neurites. Green: new neurites. **F**. Percentage of lost and new neurites in all fish (paired t-test: n=19, p=0.001). **G**. Neurite change (ratio neurite growth + ratio neurite loss) of Tph2 DRN projections in five CNS subdivisions (mixed-effects model: n(Tel/Di/Mes/Rh/SC) =6/14/17/19/10); p=0.0002). Regions with different letters are significantly different (p<0.05) after post-hoc testing. **H.** Neurite change (neurite growth +neurite loss) of Tph2 DRN projections in selected regions. DRN: Dorsal raphe nucleus, Tel: Telencephalon, Di: Diencephalon, Mes: Mesencephalon, Rh: Rhombencephalon, SC: Spinal cord, OT: Optic tectum, Dm: Dorsomedial telencephalon, A: Anterior, P: Posterior, L: Left, R: Right.

Collectively, the projections of 5-HT DRN neurons of 2 w.o. fish extended across the whole brain, innervating multiple areas in all CNS subdivisions, similar to the 1 w.o. projectome (Fig. 4B) and the overall DRN projection pattern (Sup. Fig. 4A-D). The projection profiles of selected brain regions were generally similar to those described at 1 w.o. (Sup. Fig. 4E-G). The preservation of the overall 5-HT DRN projectome appears to result from the preservation of the broad topology of individual neurons’ projection profiles, which do not undergo major changes. Indeed, neurons exhibiting either widespread or confined projection patterns largely maintained these characteristics after the second week of development (Fig. 4C, Supp. table 2), indicating that this feature is already well-established at 1 week. Interestingly, the laterality score decreased in 5-HT DRN neurons during the second week of development, suggesting a shift towards more bilateral projections (Fig. 4D, Supp. table 2).

Though no major alterations in the broad projection patterns were observed during the second week, the projections of all 5-HT DRN neurons underwent remodeling (Fig. 4E, Sup. Fig. 3). Neurite growth (27%) predominated compared to neurite loss (18%, Fig. 4F, Supp. table 2). We next asked whether these developmental changes were region-specific and found that neurite loss was uniform, whereas neurite growth differed (Sup. Fig. 4H-K, Supp. table 3). When combining growth and loss into a single measure of neurite change, the telencephalon, diencephalon, and mesencephalon showed the most pronounced remodeling (Fig. 4G, Supp. table 2), a pattern also evident at the level of individual brain regions within these subdivisions (Fig. 4H).

### Early life stress reshapes the developing 5-HT DRN projectome in a time- and region-dependent manner

To examine whether 5-HT DRN projections are sensitive to environmental changes during early life, we mapped the whole-brain density of Tph2 fibers after exposing zebrafish larvae to chronic mild stress either during the first week (Fig. 5A-D), or during the second week of life (Fig. 5E-H).

**Figure 5:**
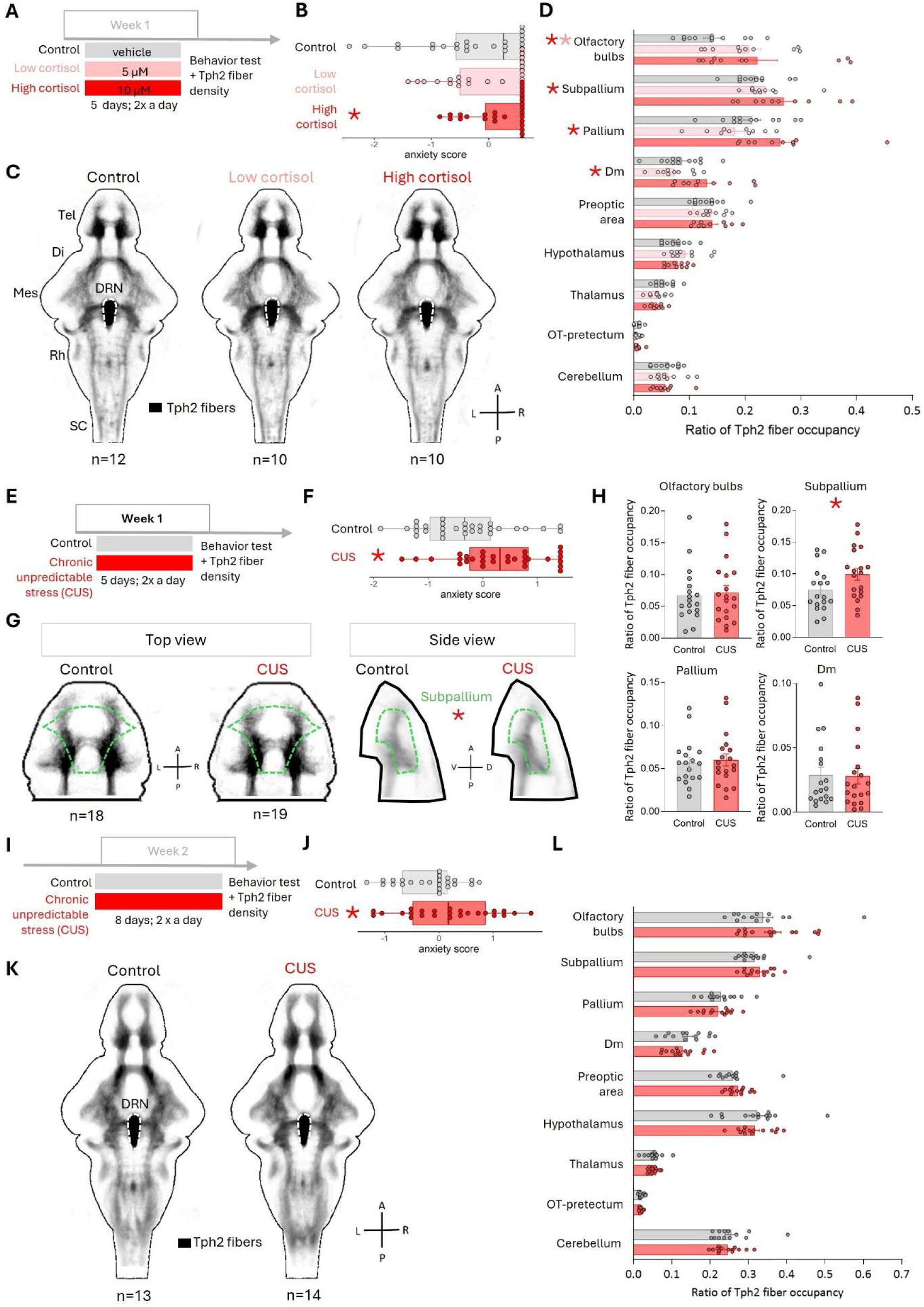
Effects of early life stress on the 5-HT DRN projectome. **A.** Scheme of the experimental design. Tg(Tph2:Gal4:UAS-caax-GFP) larval zebrafish were exposed to cortisol or vehicle twice a day from 2 to 6 days post-fertilization (control, 0 µM cortisol; low cortisol, 5 µM; high cortisol, 10 µM). Behavioral testing in the swimming plus maze and euthanasia occurred at 7 days post-fertilization. **B**. Anxiety score for all groups in the swimming plus maze assay (linear model; n=29; p(control, 5µM)=0.229; p(control, 10µM)=0.031). **C**. Z-projection maps of average Tph2 fiber density in all groups. **D**. Density of Tph2 fibers in selected CNS regions for all groups (repeated measure two-way ANOVA; n(control, low cortisol, high cortisol)=12, 10, 10; p(area:treatment)<0.0001). **E**. Scheme of the experimental design. Tg(Tph2:gal4:UAS-caax-GFP) larval zebrafish were exposed to chronic unpredictable stress (CUS) twice a day or gently handled (control) from 2 to 6 days post-fertilization. Behavioral testing in the swimming plus maze and euthanasia occurred at 7 days post-fertilization. **F.** Anxiety index in the swimming plus maze assay (one-sided student t-test, n(control,CUS)=30,30; p=0.003). **G.** Z-projection maps of average Tph2 fiber density in all groups. **H**. Density of Tph2 fibers in olfactory bulbs (one-sided student t-test, p=0.386), subpallium (p=0.027), pallium (p=0.373), and Dm (p=0.456); n(control,CUS)=18,19. **I**. Scheme of the experimental design. Tg(Tph2:gal4:UAS-caax-GFP) larval zebrafish were exposed to chronic unpredictable stress (CUS group) twice a day or gently handled (control group) from 6 to 13 days post-fertilization. Behavioral testing in the swimming plus maze and euthanasia occurred at 14 days post-fertilization. **J.** Anxiety index in the swimming plus maze assay (one-sided student t-test, n(control,CUS)=24,25, p=0.039). **K.** Z-projection maps of average Tph2 fiber density in all groups. **L**. Density of Tph2 fibers in selected CNS regions for all groups (repeated measure two-way ANOVA, n(control, CUS)=13, 14; p(area:stress)=0.597). DRN: Dorsal raphe nucleus, OT: optic tectum, Dm: Dorsomedial telencephalon, A: Anterior, P: Posterior, L: Left, R: Right. *, p<0.05 indicated for post-hoc comparisons of treated groups to controls.

Zebrafish larvae expressing GFP in broad subsets of Tph2 DRN neurons were exposed to vehicle, or to low, or high concentrations of glucocorticoids in the bath, twice a day during their first week of age (Fig. 5A). To determine whether glucocorticoid treatment altered larval behavior, we measured anxiety-like responses in the swimming plus maze test ^28^ (Fig. 5B, Sup. Fig. 5B-D). High, but not low, concentration of cortisol increased anxiety (Fig. 5B, Supp. table 4), whereas swimming speed remained similar across groups (Sup. Fig. 5A, Supp. table 4).

Tph2 fiber density was mapped throughout the brain in all groups (Fig. 5C). The 5-HT DRN projectome appears overall conserved in glucocorticoid-treated fish compared to control fish, in line with the normal larval development and intact locomotive behavior observed across groups. Comparing fiber densities across treatment groups in selected brain regions revealed a significant interaction between cortisol treatment and brain regions (Supp. table 5). Fiber density increased in the olfactory bulbs in response to the low concentration of cortisol; and in all telencephalic areas examined (olfactory bulbs, pallium, subpallium, and Dm) in response to the highest cortisol concentration (Fig. 5D). The size of the Tph2+ DRN area was unchanged in control and cortisol-treated groups (Sup. Fig. 6A), indicating that the increase in telencephalic fiber density reflects enhanced region-specific projections of pre-existing neurons and not a change in the number of Tph2+ DRN cells.

To determine whether this increase in telencephalic 5-HT DRN fibers was specific to glucocorticoid treatment or generalized across stress paradigms, we exposed larvae to chronic unpredictable stressors during the same developmental period (Fig. 5E), following a protocol established in our lab ^27,29^. Fish exposed to chronic unpredictable stress (CUS) displayed increased anxiety (Fig. 5F, Sup. Fig. 6 F-H, Supp. table 4) and reduced velocity (Sup. Fig 6E) in the swimming plus maze. Quantification of fiber density across telencephalic regions revealed a selective increase in the subpallium, whereas fiber density in the olfactory bulbs, pallium, and Dm remained unchanged (Fig. 5H, Supp. table 5). The number of Tph2+ DRN cells was similar across groups (Sup. Fig. 6B), indicating that the increased subpallial fiber density did not result from changes in the number of Tph2+ DRN neurons following CUS. Subpallium Tph2 fiber density was therefore sensitive to both stress modalities tested during week 1, whereas the effect on olfactory bulb, pallium, and Dm fiber density was specific to glucocorticoid exposure.

To determine whether Tph2 projection density remains sensitive to environmental stress beyond the first week of development, zebrafish larvae were exposed to CUS during the second week (Fig. 5I). CUS treatment increased anxiety in 2-week-old fish without affecting velocity in the swimming plus maze (Fig. 5J, Sup. Fig. 5I-L, Supp. table 4). In contrast to the first-week CUS exposure, comparing fiber densities between control and CUS-exposed subjects in selected brain regions revealed no treatment effect (Fig. 5L, Supp. table 5). Similarly, quantification of the Tph2+ DRN area revealed no significant differences between groups (Sup. Fig. 6C). Therefore, CUS exposure during the second week of life increased anxiety-like behavior without producing detectable changes in 5-HT DRN projection density in the brain areas examined.

Together, these findings show that prolonged stress remodels telencephalic serotonergic projections selectively during the first week of development, with the pattern of circuit remodeling depending on stressor modality.

## DISCUSSION

By integrating single-neuron arborization tracing, whole-brain registration, and longitudinal imaging, we constructed a comprehensive map of the whole-brain projectome for 81 individual serotonergic neurons in the DRN of young zebrafish, tracked across developmental stages.

Our analysis of DRN projectome characterization and organization revealed that 5-HT DRN neurons exhibit diverse projection patterns, featuring both confined-projecting and widespread-projecting subpopulations. Notably, we identified two distinct projection subtypes: one targeting the telencephalon and the other the spinal cord. Additionally, we observed a degree of topographic organization, with anterior DRN neurons preferentially projecting to ventral regions, such as the hypothalamus. In terms of developmental dynamics, although most projections underwent local remodeling between weeks 1 and 2, the broad projection topology was preserved.

Furthermore, we found that early life stress during the first week, but not the second week, disrupted the developmental trajectory of the 5-HT DRN projectome in the telencephalon. Together, these results provide unprecedented insight into the organization, development, and plasticity of brain-wide connectivity patterns of serotonergic neurons at single-cell resolution within a vertebrate brain.

### General features of the serotonergic system projectome

As a few studies have explored the projection pattern of individual serotonergic neurons, we examine how our findings complement and contrast with those earlier observations. A recent study reconstructed the whole brain projectome of individual serotonergic neurons in the larval zebrafish brain, including 90 5-HT DRN neurons ^30^. While some general features agree between our two studies, such as the high heterogeneity in the reach of axonal projections and the presence of both ipsilateral- and bilateral-projecting neurons, our dataset provides carefully curated registration to the template brain, with axonal projections realistically confined within the animal’s skull. In mice, a pioneer study leveraged genetic intersectional strategies to sparsely label and reconstruct the whole-brain projections of single serotonergic neurons ^10^. The authors also found that serotonergic neurons display highly heterogeneous processes, in terms of length and the brain regions they innervate. In this study, the reconstructed neurons accounted for fewer than 0.6% of the entire 5-HT population in the DRN. In contrast, our study in zebrafish comprising 81 neurons at 1 week old and 21 neurons at 2-week-old, provides a representative sample of the 5-HT DRN population at both developmental stages, thereby reinforcing the robustness of our results.

Consistent with earlier work ^13^, our results confirm that the spinal cord of young zebrafish receives substantial input from 5-HT DRN neurons. This pattern contrasts with observations in rodents, where DRN projections primarily target the forebrain from embryonic stages through adulthood ^12,31^, and where less than 3% of spinal motor neuron innervation originates from the DRN^32^. Interestingly, this difference may be age-dependent, as it has been proposed that serotonergic projections from the rostral raphe to the spinal cord does not persist into adulthood in zebrafish ^13^. Our discovery of two broad subsets of neurons, one projecting to the telencephalon and the other to the spinal cord during the first two weeks of development, represents a previously unreported organizational feature of the DRN and may help resolve the apparent discrepancy between larval and adult 5-HT DRN projectomes. Specifically, this organization in the larva could allow for the selective pruning or apoptosis of spinal cord-projecting neurons later in development, while preserving neurons innervating other brain regions. Taken together, our findings and the existing literature point to a species-specific process in which the DRN modulates locomotion in young zebrafish via direct serotonergic projections to the spinal cord.

Another insight gained from our study is that most neurons arbors bilateral projections, with 58% of 5-HT DRN neurons displaying projections to both ipsilateral and contralateral hemispheres at 1 week old. Furthermore, we observed an increase in the proportion of bilateral projections by 2 weeks of age. These results suggest that the serotonergic system in young zebrafish likely engages in extensive synchronous modulation across both brain hemispheres. Examples of functions involving the serotonergic system that might require coordinated action across hemispheres include sleep ^33^ and sensory processing ^34^.

### Developmental remodeling of 5-HT DRN projections

In mammals, serotonergic projections are established during embryonic and early postnatal development, with a region-specific timing ^12,16,35^. In the present study, we also found region-specific changes in the density 5-HT fibers in the zebrafish CNS. For example, processes to the anterior portion of the spinal cord are already established by 1 week and do not grow further during the second week. This finding highlights the early formation and stability of 5-HT modulation in spinal circuits, aligning with the onset of swimming behavior at 4–5 dpf ^36^. Furthermore, we measured high levels of remodeling in 5-HT DRN fibers during the second week in the preoptic area, the hypothalamus and the pallium. These brain regions are known to mediate a range of functions, including social preference ^37,38^ and associative learning, both of which develop from 2-3 week of age ^39^. Together, our results suggest that the timing of innervation of these areas by the 5-HT DRN may coincide with the functional maturation of specific neural circuits, thereby supporting the emergence of behavioral and cognitive abilities during early development.

Tracking the same neuron over time through longitudinal imaging is a valuable method for uncovering dynamic changes in neuronal processes during development. However, such studies are particularly challenging in mammals, where only neurons with local projections and limited spatial reach can be reliably reconstructed and monitored over several days in the living brain. Hence, this approach has been primarily used in insects or in small vertebrate models ^40,41^. In zebrafish, time-lapse imaging has been used to visualize the processes of individual sensory neurons, sometimes over several days and up to 8 days post-fertilization ^42–44^.

To our knowledge, our study is the first to reconstruct the complete axonal arbor of single neuromodulatory neurons across the entire brain and to compare their structure at one-week interval during vertebrate development. The use of membrane-tagged GFP in our study allowed accurate visualization and tracking of the entire neurite tree and revealed novel principles of plasticity within the developing 5-HT system. Insights in the dynamic changes in 5-HT innervation during development come from immunolabelling, or from transgenic mice expressing GFP broadly across Tph2 neurons ^12,16,35^. Therefore, it is unknown whether changes in fiber density are driven by global modification of all 5-HT neurons’ processes, or by changes in only a subset of 5-HT neurons. We found that all neurons exhibited some degree of neurite remodeling during the second week of development, indicating that morphological plasticity is not restricted to specific neurons but is instead a shared property of the 5-HT DRN population at this developmental stage. However, the extent of neurite pruning and growth varied considerably among neurons, showing that their projections do not remodel to the same degree. Despite this widespread structural plasticity, the broad projection topology was preservedsuggesting that the main features of the 5-HT DRN projectome are well-established and functionally relevant by one week of age.

### A critical window of sensitivity to early-life stress

We show that both cortisol treatment and chronic unpredictable stress exposure during the first week of life (days 2-6) altered 5-HT DRN fiber density in a region-specific manner, whereas chronic unpredictable stress during the second week did not. These findings suggest that serotonergic projections are particularly sensitive to environmental perturbations during the first week of development, when most projection patterns of 5-HT neurons are being established. This is consistent with previous work identifying a critical window between days 2 and 6, during which increased sensitivity to early-life stress significantly impacts the later development of anxiety-like symptoms ^45^. Our results extend these findings by suggesting that this critical window also applies to the development of 5-HT DRN projections. Interestingly, cortisol and CUS produced distinct patterns of remodeling, with cortisol inducing broader changes across telencephalic regions and CUS selectively increasing fiber density in the subpallium. These differences suggest that developmental timing defines a period of heightened susceptibility, whereas the specific remodeling pattern might depend on stressor modality.Interestingly, the telencephalic regions affected by cortisol are among those where 5-HT fibers grows most during the second week of life (Sup. Fig. 5K). Our results suggest that cortisol exposure during the first week might accelerate the maturation of 5-HT projections into one resembling a 2 week-like pattern. Accelerated maturation of connectivity between brain regions, particularly within the corticolimbic system, has been well documented in children and animal models exposed to early life trauma ^46,47^.

Our findings do not support a simple relationship between alterations in DRN serotonergic projections and anxiety-like behavior. Across three experiments in which environmental stress levels were manipulated during distinct developmental periods, all stress paradigms increased anxiety-like behavior, whereas changes in serotonergic fiber density were observed in only two of the three conditions. These results suggest that chronic stress can promote anxiety-like behavior without necessarily inducing large-scale remodeling of serotonergic projections. Consistent with this interpretation, previous work from our group showed that chronic unpredictable stress affects the activity of DRN serotonergic neurons and contributes to altered habituation, but not to anxiety-like behavior^27^.

In conclusion, our work reveals principles of projectome establishment and maintenance for an important neuromodulatory hub. It provides a resource for future studies of neuronal development and of the organizational logic of sparse neuromodulatory systems.

## SUPPLEMENTARY INFORMATION

**Supplementary figure 1:**
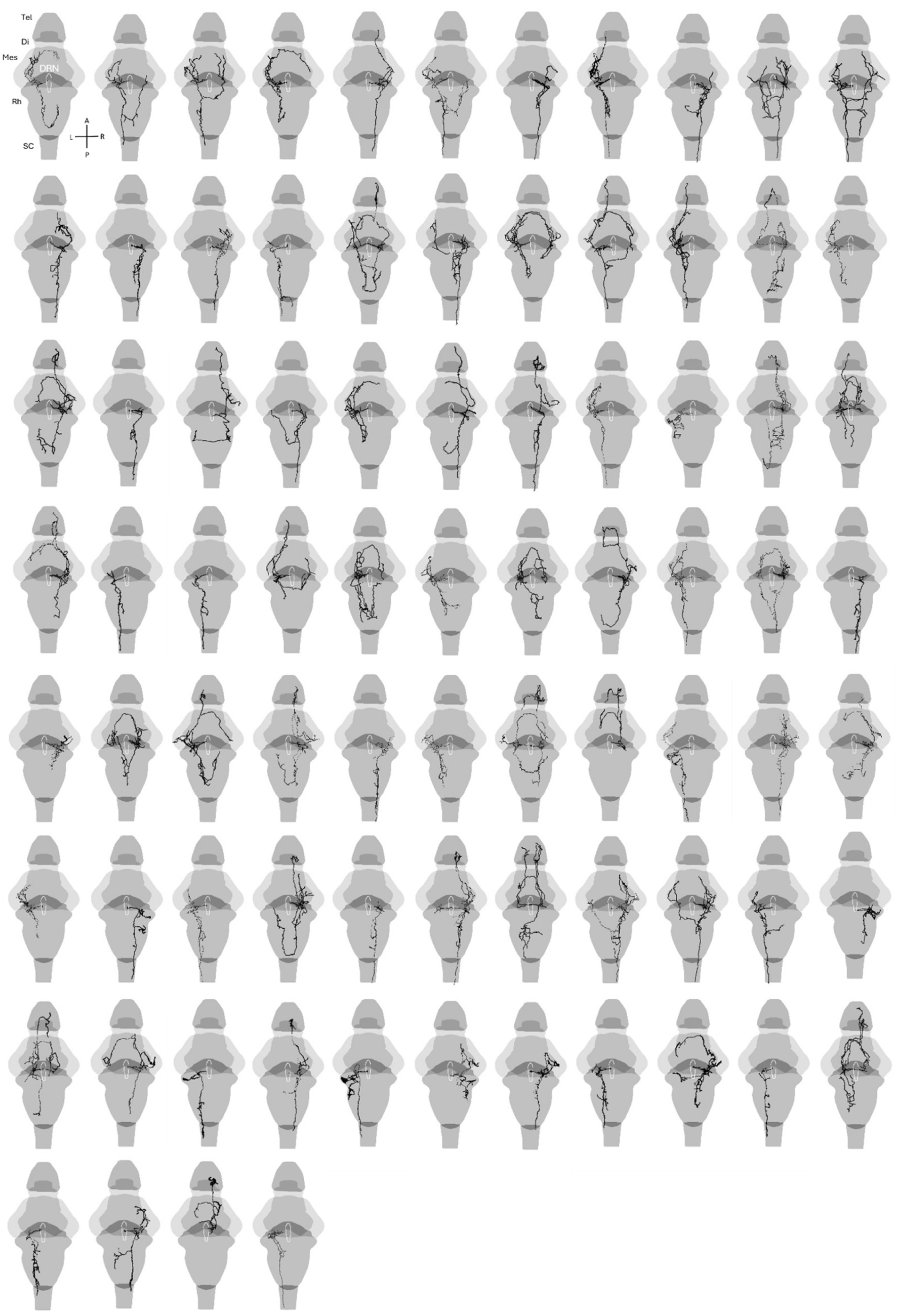
Whole-brain projections of 81 5-HT DRN neurons in 1 week-old larvae. Depth projection of all individually reconstructed Tph2 neurons. DRN: Dorsal raphe nucleus, Tel: Telencephalon, Di: Diencephalon, Mes: Mesencephalon, Rh: Rhombencephalon, SC: Spinal cord, A: Anterior, P: Posterior, L: Left, R: Right.

**Supplementary figure 2:**
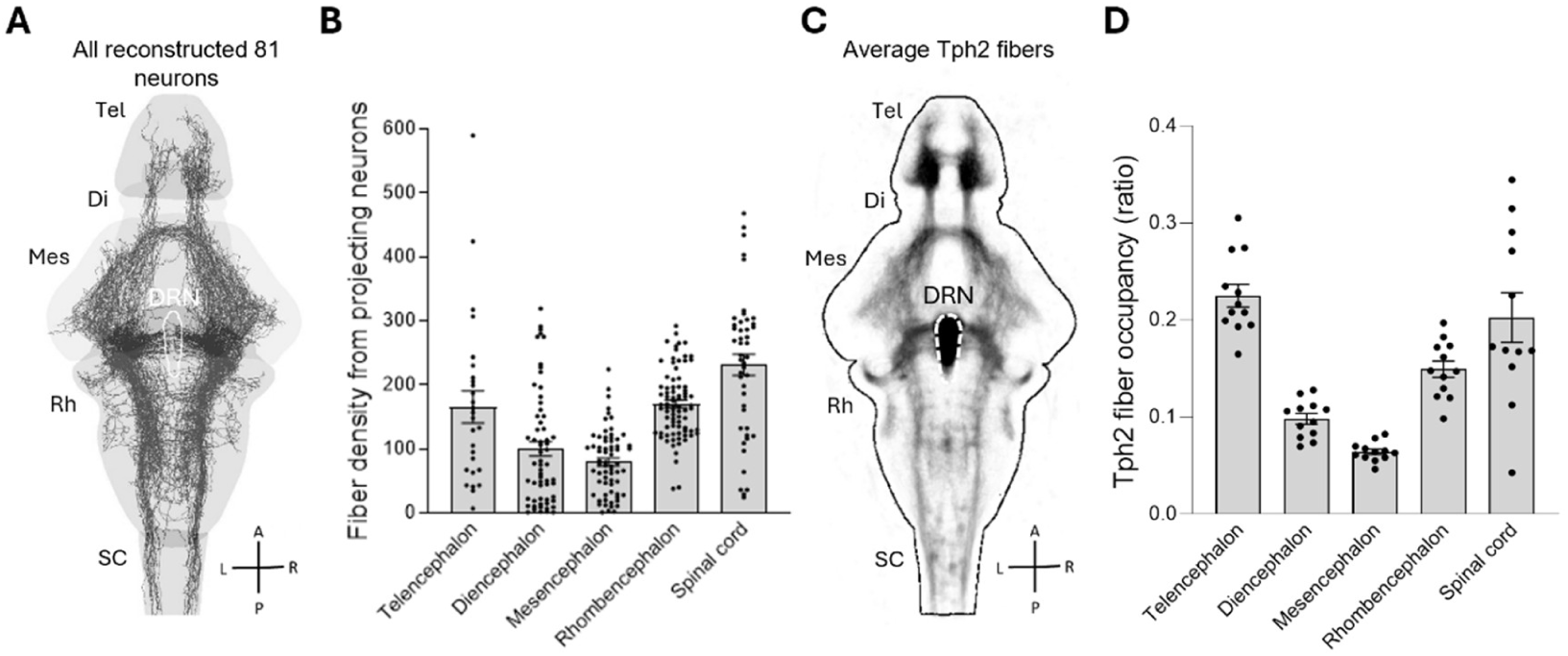
Whole-brain Tph2 fiber density in sparsely and broadly labelled fish. **A**. Depth projection of all individually reconstructed Tph2 neurons at 1 week old (n=81 neurons). **B**. Tph2 fiber density in main CNS subdivisions corresponding to data in A. Density = 4.10^-9^ fiber’s area/ brain region area. **C**. Average depth projection of Tph2 fibers in 1 week old fish with broadly labelled dorsal raphe nucleus (n=12 fish). **D**. Tph2 fiber density in main CNS subdivisions corresponding to data in C. DRN: Dorsal raphe nucleus, Tel: Telencephalon, Di: Diencephalon, Mes: Mesencephalon, Rh: Rhombencephalon, SC: Spinal cord, A: Anterior, P: Posterior, L: Left, R: Right.

**Supplementary figure 3:**
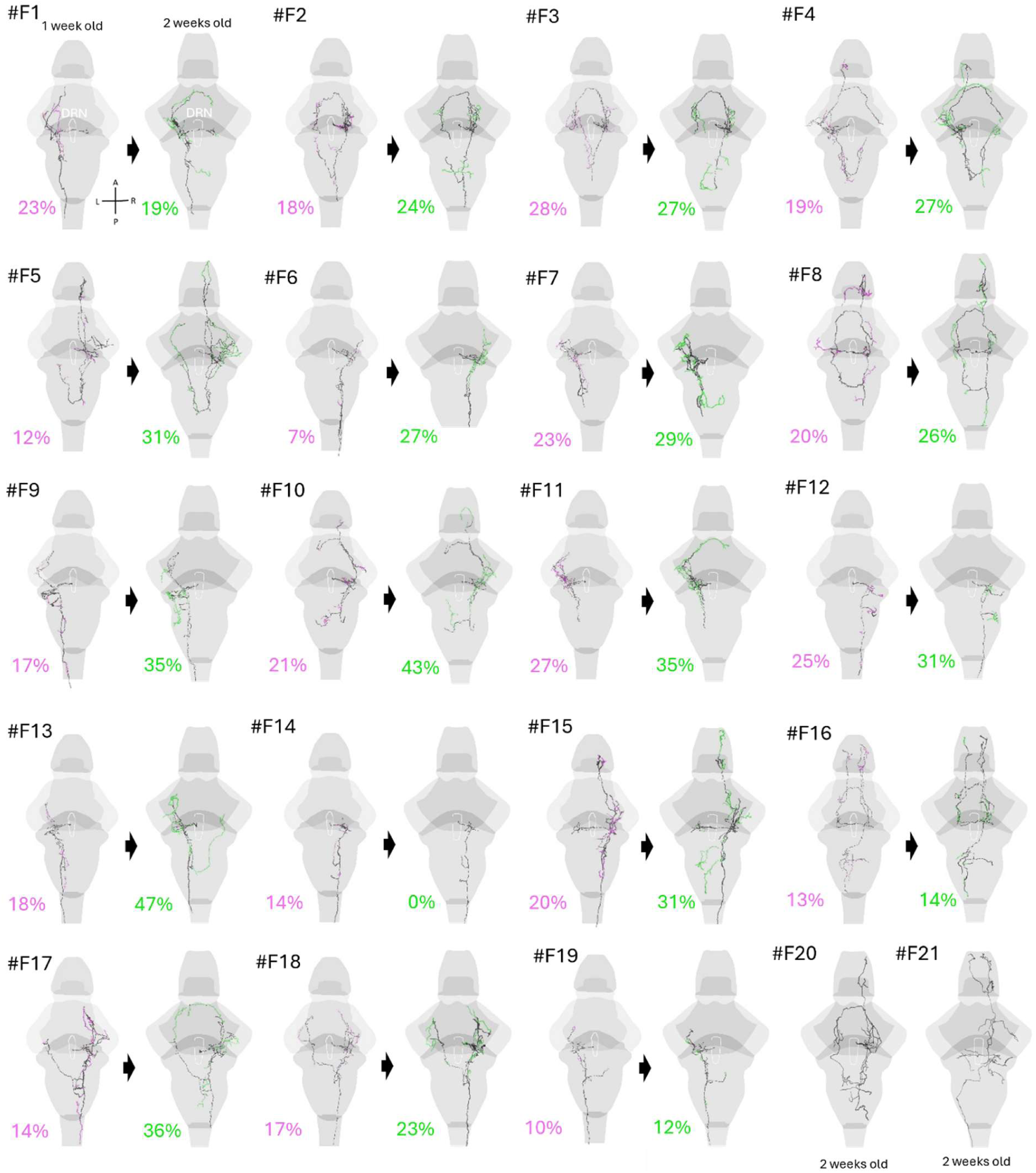
Projections of longitudinally imaged 5-HT DRN neurons. Depth projection of reconstructed Tph2 neurons imaged at both the 1-week-old and the 2-week-old time points. Magenta colored traces and percentages correspond to pruned neurites. Green-colored traces and percentages correspond to new neurites. Fish 6 (#F6) was excluded from the analysis involving the spinal cord due to missing this area in the 2 w.o. scan. #F20 and #F21 are included only in the analyses of the overall 2 w.o. projectome. DRN: Dorsal raphe nucleus, A: Anterior, P: Posterior, L: Left, R: Right

**Supplementary figure 4:**
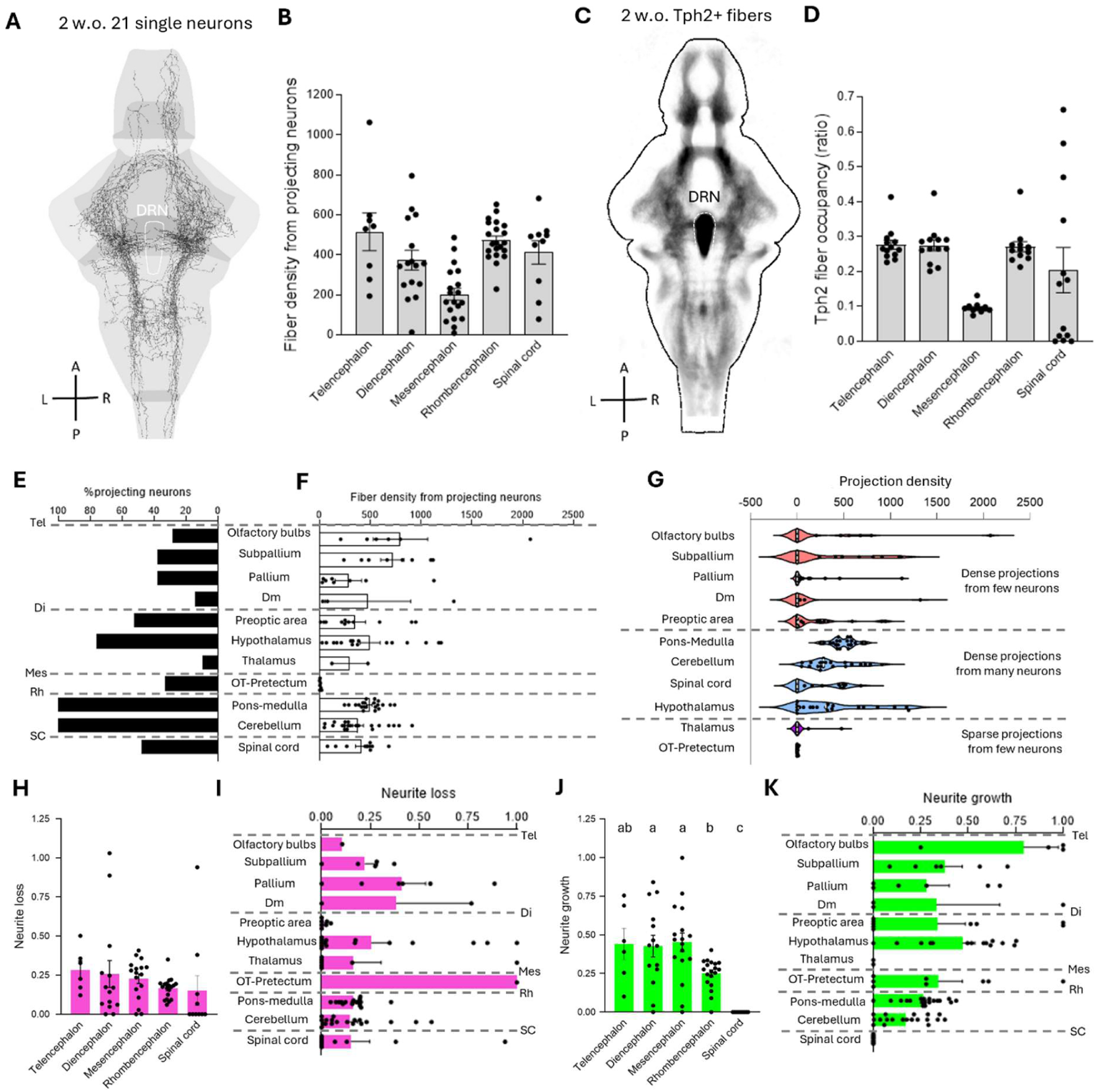
Longitudinal imaging of the developing 5-HT DRN projectome. **A**. Overlay of all Tph2 neurons at 2 weeks old (n=21 neurons). **B**. Tph2 fiber density in main CNS subdivisions corresponding to data in A. Density = 4.10^-9^ fiber’s area/ brain region area. **C**. Average projection of Tph2 fibers in 2 weeks old fish with broadly labelled dorsal raphe nucleus (n=13 fish). **D**. Tph2 fiber occupancy in main CNS subdivisions corresponding to data in C. **E.** Percentage of reconstructed 2 w.o. Tph2 neurons projecting to diverse brain regions. **F.** Density of Tph2 fibers from projecting neurons at 2 w.o. in diverse brain regions (Mean ±SEM). Density = 4.10^-9^ fiber’s area/ brain region area. **G.** Distribution of projection density by all 2 w.o. Tph2 neurons across diverse brain regions, distinguishing the three categories of regions identified by the zero-inflated model. Density = 4.10^-9^ fiber’s area/ brain region area. **H**. Ratio of Tph2 fiber loss in the five CNS subdivisions (mixed-effects model: number neurons, n(Tel,Di,Mes,Rh,SC)=6,14,17,19,10; p=0.503). Neurite loss=density lost trace/density whole trace. **I.** Ratio of Tph2 fiber loss in selected brain regions (Mean ±SEM). **J**. Ratio of Tph2 fiber growth in the five CNS subdivisions (mixed-effects model: number neurons, n(Tel,Di,Mes,Rh,SC)=6,14,17,19,10; p<0.0001). Regions with different letters are significantly different (p<0.05) after post-hoc testing. Neurite growth=density new trace/density whole trace. **K.** Ratio of Tph2 fiber growth in selected brain regions (Mean ±SEM). DRN: dorsal raphe nucleus, Tel: telencephalon, Di: Diencephalon, Mes: Mesencephalon, Rh: Rhombencephalon, SC: Spinal cord, OT: Optic tectum, Dm: Dorsomedial telencephalon. A: Anterior, P: Posterior, L: Left, R: Right.

**Supplementary figure 5:**
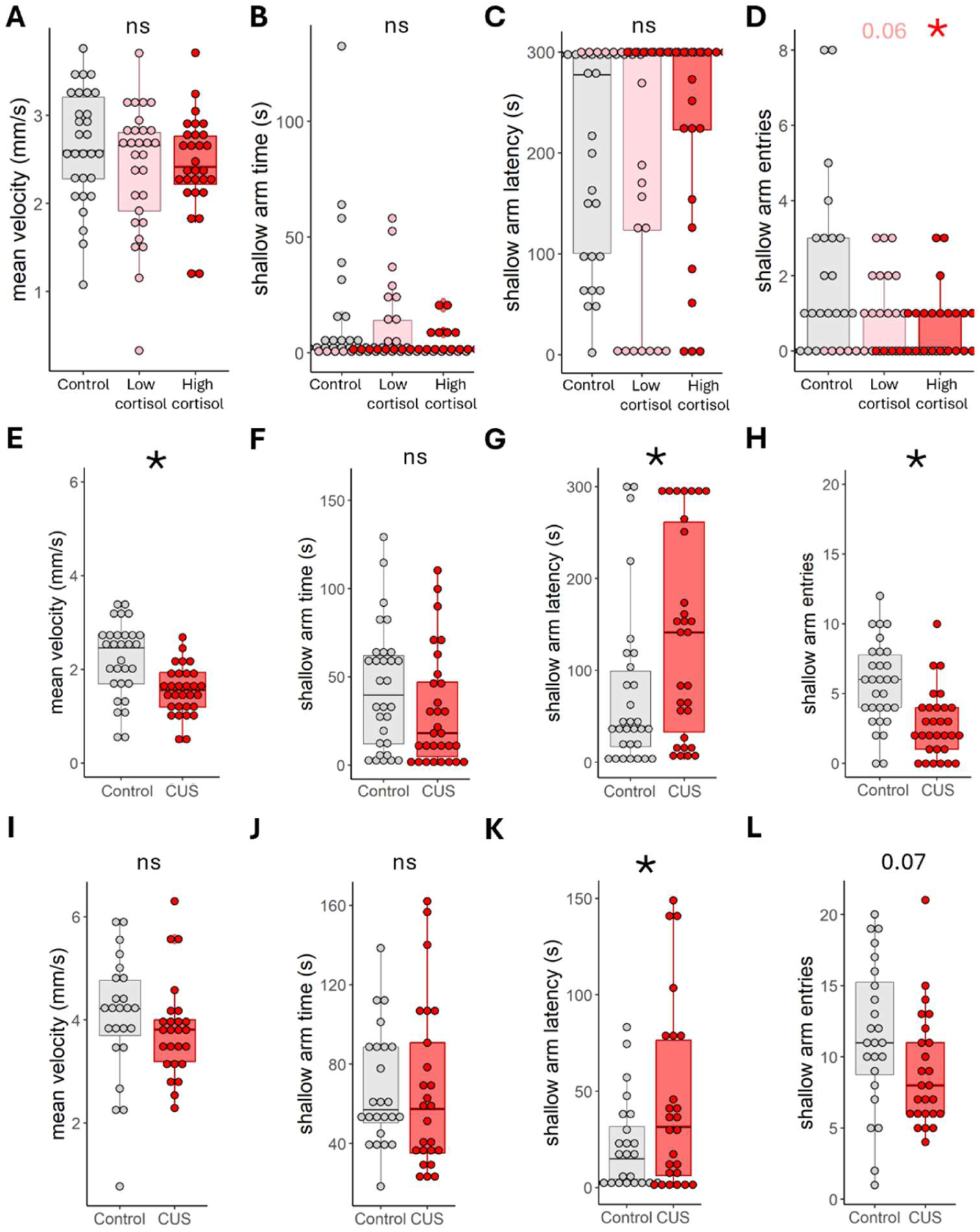
Effects of chronic early life stress on locomotive and anxiety-like behaviors in the swimming plus maze. **A.** Mean velocity of cortisol-exposed and control fish (linear model; n=29; p=0.246). **B.** Time spent in shallow arms by cortisol-exposed and control fish (linear model; n =29 in all groups; p(control vs. 5µM)=0.589; p(control vs. 10 µM)=0.069). **C.** Latency to first entry to shallow arms of cortisol-exposed and control fish (linear model; p=0.351). **D.** Number of entries to shallow arms of cortisol-exposed and control fish (linear model; n=29; p(control vs. 5µM)=0.062; p(control vs. 10µM)=0.018). **E.** Mean velocity of 1 w.o. CUS-exposed and control fish (student t-test; n=30 per group; p<0.001). **F.** Time spent in shallow arms by 1 w.o. CUS-exposed and control fish (p=0.126). **G.** Latency to first entry to shallow arm of 1 w.o. CUS-exposed and control fish (p=0.016). **H**. Number of entries to shallow arm of 1 w.o. CUS-exposed and control fish (p<0.001). **I.** Mean velocity of 2 w.o. CUS-exposed and control fish (student t-test; n(control,CUS)=24,25; p=0.533). **J.** Time spent in shallow arms by 2 w.o. CUS-exposed and control fish (p=0.954). **K.** Latency to first entry to shallow arm of 2 w.o. CUS-exposed and control fish (p=0.048). **L**. Number of entries to shallow arm of 2 w.o. CUS-exposed and control fish (p=0.068).

**Supplementary figure 6:**
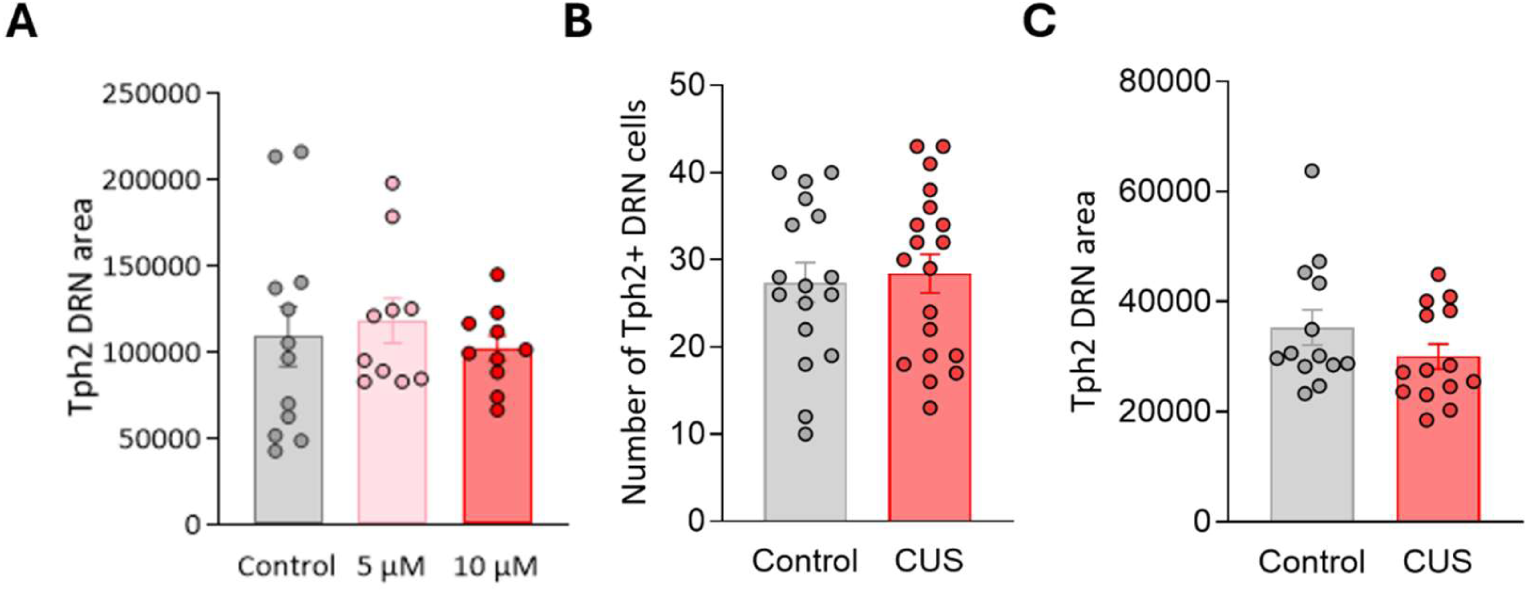
Quantification of the tph2+ dorsal raphe nucleus area after chronic early life stress. **A.** Mean size (in pixels) of tph2+ DRN of cortisol-exposed and control fish (one-way ANOVA; n(control, low cortisol, high cortisol)=12, 10, 10; p=0.422). **B.** Number of tph2+ DRN soma in 1 w.o. CUS-exposed and control fish (student t-test; n(control,CUS)=17,19; p=0.757). One close-up scan of the DRN was missing, otherwise same fish as those in Figure 5 G-H. **C.** Mean size (in pixels) of tph2+ DRN of 2 w.o. CUS-exposed and control fish (student t-test; n(control,CUS)=13,14; p=0.189).

**Table 1:**
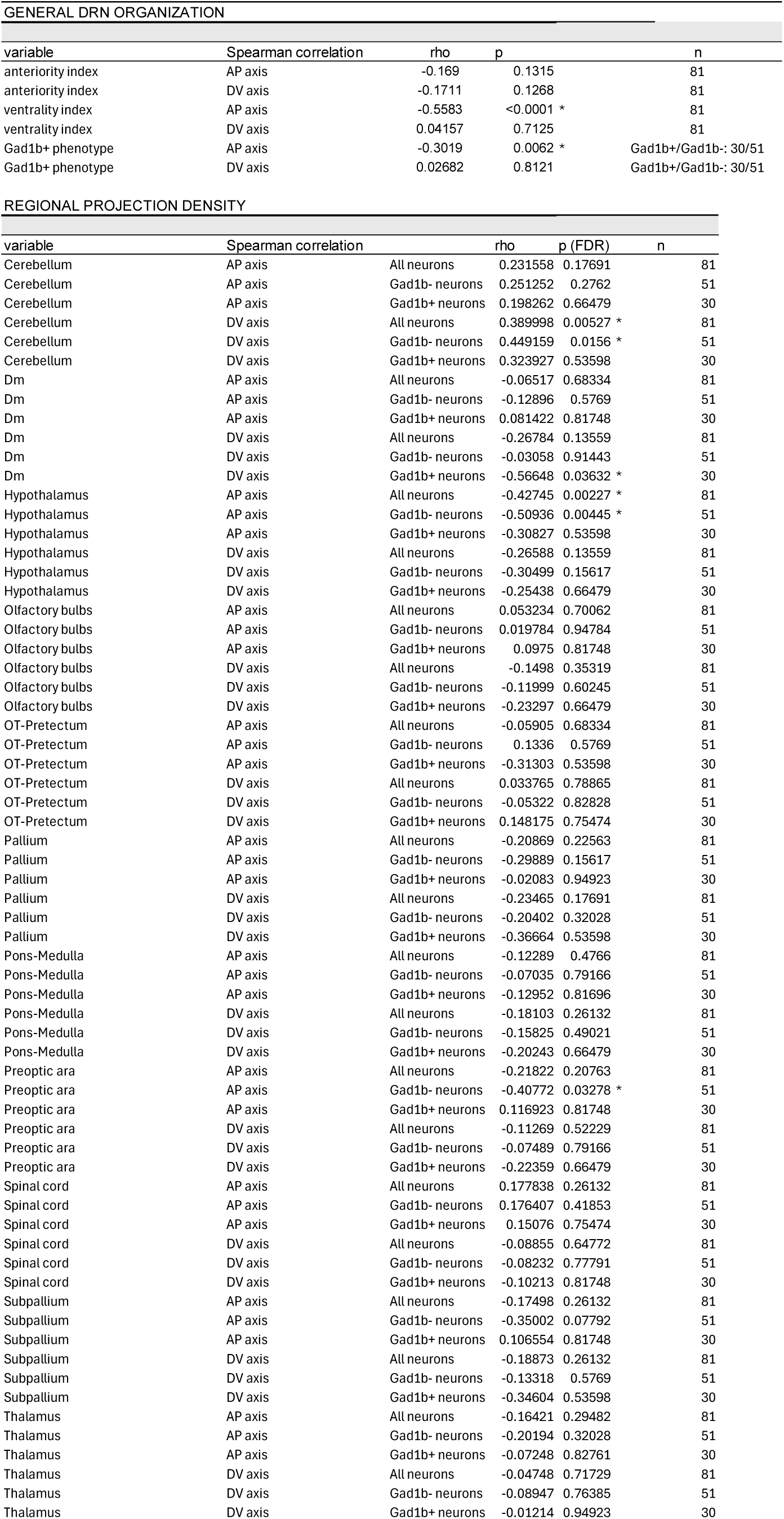
Topographical organization of 5-HT DRN neurons’ projections.

**Table 2:** Development of 5-HT DRN neurons’ projections.

| Table 2: Development of 5-HT DRN neurons' projections |  |  |  |  |  |  |
| --- | --- | --- | --- | --- | --- | --- |
| CHANGE IN PROJECTION SCORE |  |  |  |  |  |  |
| 1 w.o. vs. 2 w.o. |  | one animal was excluded: missing spinal cord in 2 w.o. scan |  |  |  |  |
| variable | paired t.test | effect size ($\eta^2$) | df | t | p | n |
| projection score 1 w.o. | projection score 2 w.o. | 0.056 | 17 | 1 | 0.3313 | 18 |
| CHANGE IN LATERALITY SCORE |  |  |  |  |  |  |
| 1 w.o. vs. 2 w.o. |  |  |  |  |  |  |
| variable | paired t.test | effect size ($\eta^2$) | df | t | p | n |
| laterality score 1 w.o. | laterality score 2 w.o. | 0.263 | 18 | 2.531 | 0.0209 * | 19 |
| OVERALL NEURITE GROWTH AND LOSS |  |  |  |  |  |  |
| % neurite loss vs. %new neurites |  |  |  |  |  |  |
| variable | paired t.test | effect size ($\eta^2$) | df | t | p | n |
| % neurite loss | %new neurites | 0.440 | 18 | 3.758 | 0.0014 * | 19 |
| NEURITE CHANGE |  |  |  |  |  |  |
| variable | Mixed-effects model (REML) | effect size (partial $\eta^2$) | df | F | p | n |
| ratio neurite change | area | 0.390 | 4, 43 | 6.862 | 0.0002 * | Tel/Di/Mes/Rh/SC: 6/14/17/19/10 |
| variable | posthoc | effect size (cohens d) | df | t | p (FDR) | n |
| ratio neurite change |  |  |  |  |  |  |
| Telencephalon | Diencephalon | 0.103 |  | 0.1266 | 0.987 | Tel/Di: 6/14 |
| Telencephalon | Mesencephalon | 0.174 |  | 0.1425 | 0.987 | Tel/Mes: 6/17 |
| Telencephalon | Rhombencephalon | 1.745 |  | 1.976 | 0.078 | Tel/Rh: 6/19 |
| Telencephalon | Spinal cord | 2.212 |  | 3.45 | 0.0042 * | Tel/SC: 6/10 |
| Diencephalon | Mesencephalon | 0.015 |  | 0.0164 | 0.987 | Di/Mes: 14/17 |
| Diencephalon | Rhombencephalon | 0.711 |  | 2.512 | 0.0317 * | Di/Rh: 14/19 |
| Diencephalon | Spinal cord | 1.260 |  | 4.205 | 0.0006 * | Di/SC: 14/10 |
| Mesencephalon | Rhombencephalon | 1.070 |  | 2.651 | 0.028 * | Mes/Rh: 17/19 |
| Mesencephalon | Spinal cord | 1.699 |  | 4.357 | 0.0006 * | Mes/SC: 17/10 |
| Rhombencephalon | Spinal cord | 1.124 |  | 2.255 | 0.0488 * | Rh/SC: 19/10 |

**Table 3:** Neurite loss and growth of individual 5-HT DRN neurons.

| Table 3: Neurite loss and growth of individual 5-HT DRN neurons |  |  |  |  |  |  |
| --- | --- | --- | --- | --- | --- | --- |
| NEURITE LOSS |  |  |  |  |  |  |
| variable | Mixed-effects model (REML) | effect size (partial $\eta^2$) | df | F | p | n |
| ratio neurite loss | area | 0.073 | 4, 43 | 0.8466 | 0.5035 | Tel/Di/Mes/Rh/SC: 6/14/17/19/10 |
| NEURITE GROWTH |  |  |  |  |  |  |
| variable | Mixed-effects model (REML) | effect size (partial $\eta^2$) | df | F | p | n |
| ratio neurite growth | area | 0.499 | 4, 42 | 10.44 | <0.0001 * | Tel/Di/Mes/Rh/SC: 6/14/17/19/10 |
| variable | posthoc | effect size (cohens d) | df | t | p (FDR) | n |
| ratio neurite growth |  |  |  |  |  |  |
| Telencephalon | Diencephalon | 0.056 |  | 0.01369 | 0.9891 | Tel/Di: 6/14 |
| Telencephalon | Mesencephalon | -0.040 |  | 0.3082 | 0.8438 | Tel/Mes: 6/17 |
| Telencephalon | Rhombencephalon | 1.009 |  | 1.92 | 0.0882 | Tel/Rh: 6/19 |
| Telencephalon | Spinal cord | 2.493 |  | 4.19 | 0.0005 * | Tel/SC: 6/10 |
| Diencephalon | Mesencephalon | -0.095 |  | 0.397 | 0.8438 | Di/Mes: 14/17 |
| Diencephalon | Rhombencephalon | 0.896 |  | 2.622 | 0.0202 * | Di/Rh: 14/19 |
| Diencephalon | Spinal cord | 2.301 |  | 5.211 | <0.0001 * | Di/SC: 14/10 |
| Mesencephalon | Rhombencephalon | 1.064 |  | 3.212 | 0.0051 * | Mes/Rh: 17/19 |
| Mesencephalon | Spinal cord | 2.553 |  | 5.746 | <0.0001 * | Mes/SC: 17/10 |
| Rhombencephalon | Spinal cord | 3.677 |  | 3.286 | 0.0051 * | Rh/SC: 19/10 |

**Table 4:**
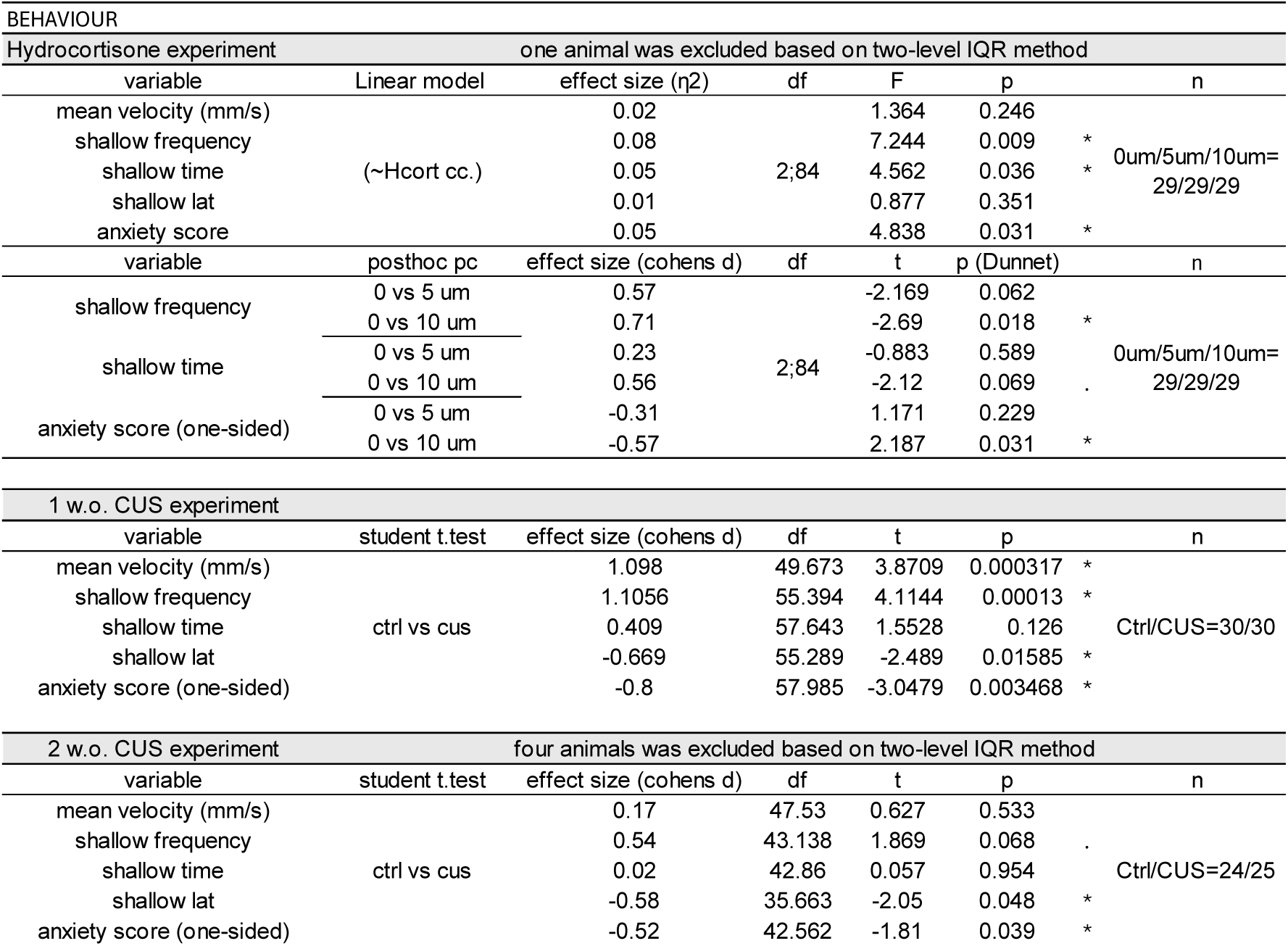
Effect of ELS on locomotion and anxiety-like behavior.

**Table 5:**
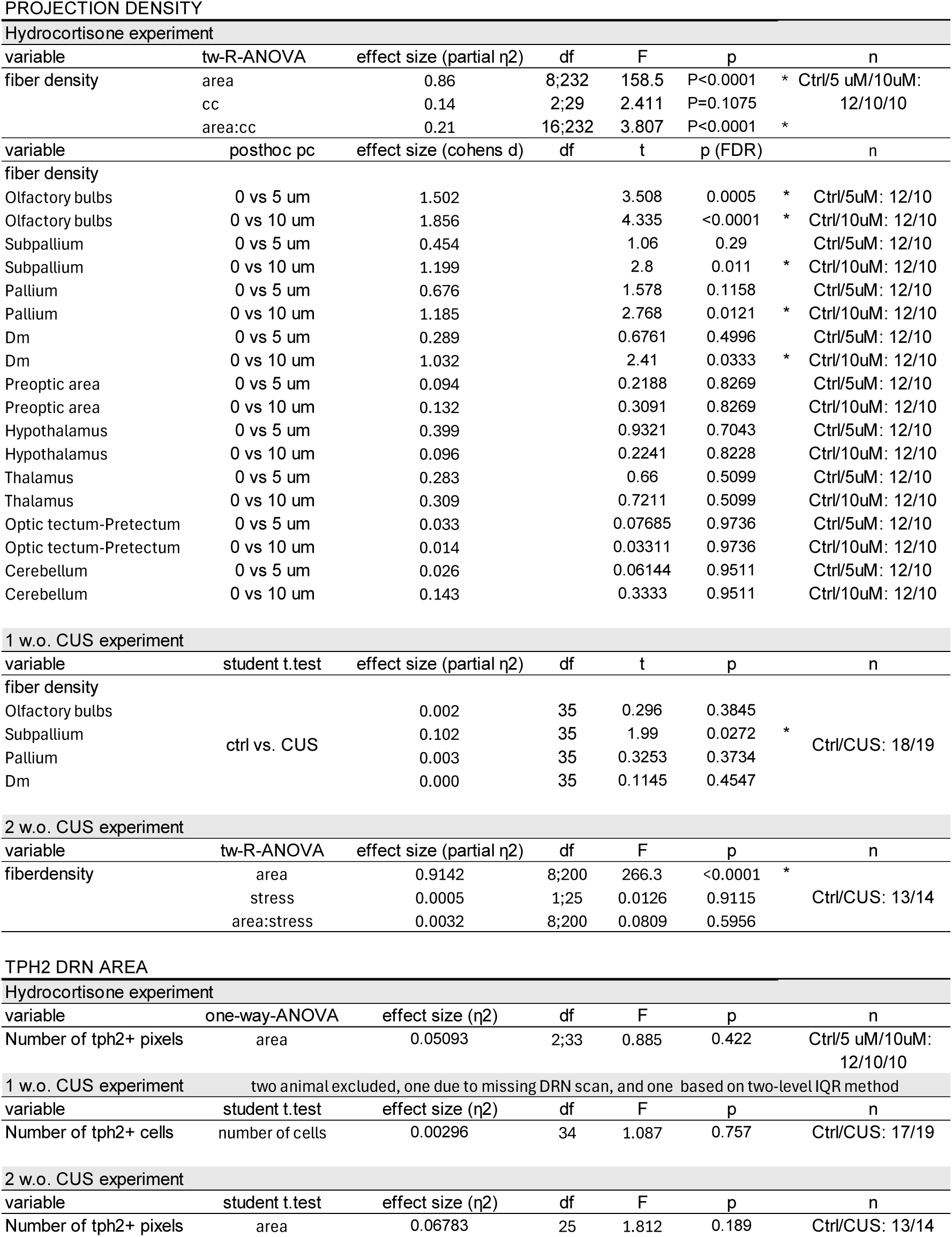
Effect of ELS on the 5-HT DRN projectome.

**Table 6:**
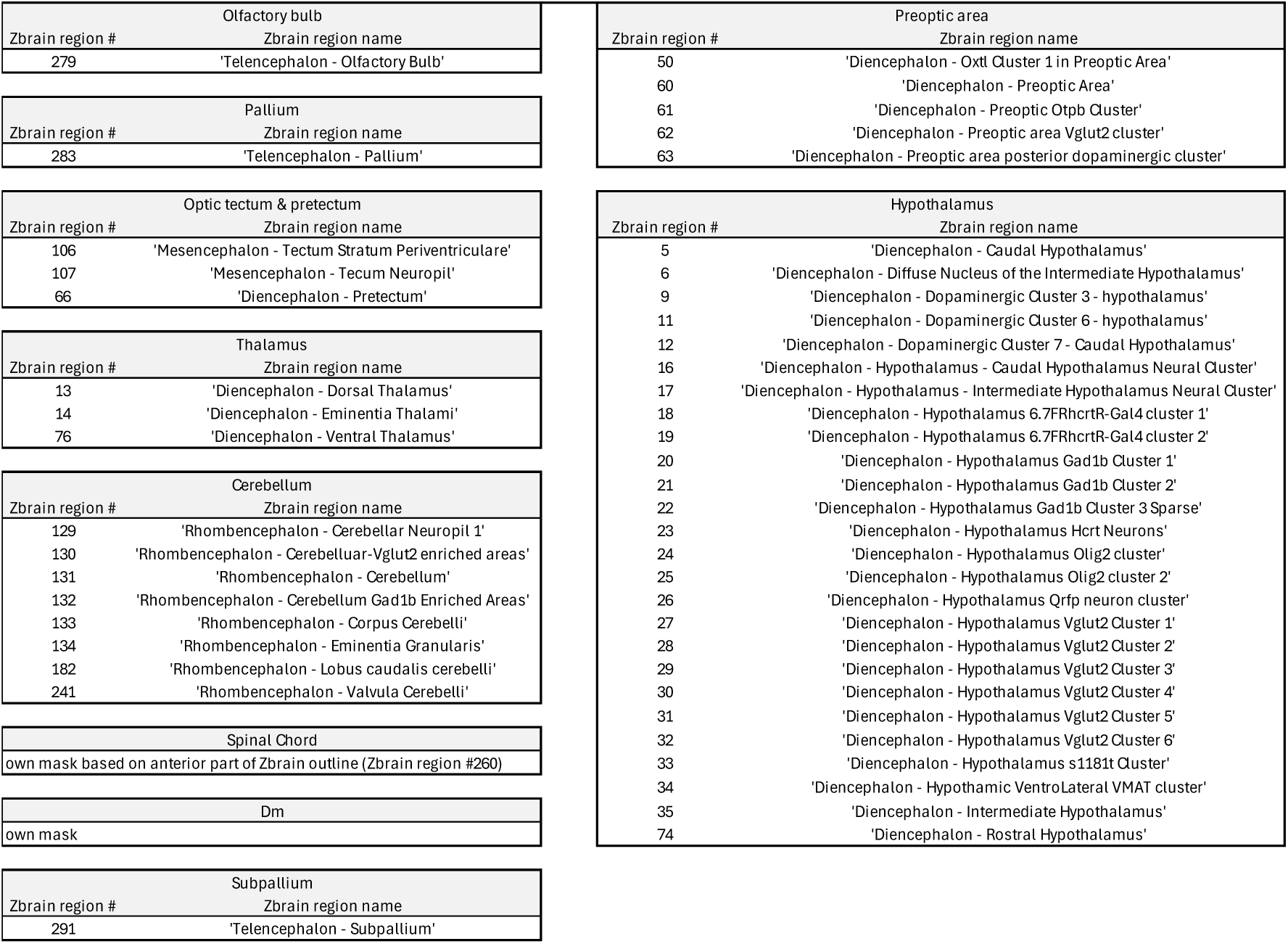
Definition of brain regions used in the study.

## METHODS

### Animals

Zebrafish were six to fourteen days old at the time of the experiments. We used the following transgenic lines, in the *nacre mifta^-/-^*background ^48^: Tg(*tph2:gal4;UAS:Caax-GFP*) expressing the membrane-tagged green fluorescent protein (GFP) under the tryptophan hydroxylase 2 promoter in serotonergic neurons of the dorsal raphe nucleus ^4^, Tg(*gad1b:DsRed*) ^49^ expressing the red fluorescent protein DsRed under the Glutamate decarboxylase 1 (Gad1) promoter, Tg(*elavl3:H2b:GCaMP6s*), Tg(*pet1:mCherry*) ^13^ and Tg(*tbx21:Gal4:UAS:tdTomato*) ^50^. Larvae were raised in petri dishes at 28°C in an incubator at a density of approximately 60 larvae/50 mL in artificial fish water (AFW; 0.3 g marine salt in 5 L deionized water). At 2 days post-fertilization (dpf), fish used in the cortisol exposure experiments were relocated to shallow-water tanks (length × width × height: 18 × 10.5 × 8 cm) equipped with meshed-bottom nursery inlets (length × width × height: 18 × 10.5 × 6.5 cm). Fish used in the chronic unpredictable stress experiments were transferred at 5 dpf to a recirculating system (Techniplast) in 3.5 L tanks with nursery inlets. In the recirculating system, animals were raised at 28.4 °C, 400 μS/cm, pH of 7.4 under a 10/14-hour light cycle. From 5dpf, fish were fed with commercial food (shrimp larval diet, Royal Caviar, 5-50 μm). All animal experiments were conducted following the guidelines of the Danish Animal Experimentation Inspectorate (permit 2023-15-0201-01493) in a fully AAALAC-accredited facility under the supervision of the local animal welfare committee.

### Experimental design

#### Single neurons’ projections experiments

Tg(*tph2:Gal4;UAS:Caax-GFP; gad1b:DsRed*) fish were screened at 4-5 dpf for DsRed expression throughout the brain and GFP expression in single neurons using an epifluorescence microscope (Leica M165 FC, Leica Microsystems). At 6 or 7 dpf, fish were anesthetized and imaged under a confocal microscope. Only 2 of the 81 imaged neurons originated from the same animal.

#### Development experiments

For longitudinal imaging of the same neuron across development, Tg(*tph2:Gal4;UAS:Caax-GFP; gad1b:DsRed*) fish expressing GFP in a single neuron were imaged twice, at 1 w.o. and 2 w.o. Following the first scan, fish were gently extracted from the agar, rinsed in AFW, and returned to the home petri dish. To avoid social isolation while retaining the identity of scanned fish, each of them was raised in a separate petri dish, together with age-matched pigmented conspecifics from the AB line. All imaged neurons originated from different animals.

#### Cortisol exposure experiments

At 2 dpf, eggs were randomly allocated to control, 5 µM cortisol, and 10 µM cortisol groups, and placed in tanks with nursery inlets that enabled easier transfer to the treatment tanks. Fish in the cortisol-treated groups were exposed to cortisol from 2 to 6 dpf. The swimming plus-maze test was performed at 7 dpf, and fish were euthanized and fixed at the end of the assays.

#### Chronic unpredictable stress experiments

Larvae were randomly assigned to control or stressed groups at 1 dpf for the 1 w.o. experiment or at 5 dpf for the 2 w.o. experiment. Fish were housed in tanks with nursery inlets. From 2 to 6 dpf (1 w.o. experiment) or from 6 to 13 dpf (2 w.o. experiment), the fish in the stressed group were subjected to the chronic unpredictable stress protocol. The swimming plus-maze test was performed at 7 dpf (1 w.o. experiment) or at 14 dpf (2 w.o. experiment), and the fish were euthanized and fixed after the assays.

### Early life stress exposure

#### Cortisol exposure

Water-soluble hydrocortisone 21-hemisuccinate sodium salt (Merck, H4881) was dissolved in AFW to a concentration of 5 µM or 10 µM. From 2 to 6 dpf, the cortisol-treated fish were bathed in the corresponding doses of cortisol twice a day for 20 minutes at random times of the day between 11 am and 6 pm. Following cortisol exposure, animals were rinsed in AFW (2 x 1min). Control fish were subjected to the same handling but treated with AFW.

#### Chronic unpredictable stress (CUS)

From 2 to 6 dpf (1 w.o. experiment) or from 6 to 13 dpf (2 w.o. experiment), the fish in the stressed group were exposed to two stressors per day, applied at random times between 8 am and 8 pm to maintain unpredictability, according to a protocol previously established in our lab ^29^. The following five stressors were used: chasing, turbulence, hyperosmotic shock (100 mM NaCl), pH drop (pH = 4), and light flashes exposure (6 mW/cm2 light flashes at 5 Hz). Fish in the control group remained undisturbed except for gentle tank transfer twice a day. The animals received additional feeding 20-30 minutes after exposure to the stressors or tank transfer.

### Swimming plus maze

For all behavioral tests, fish were transferred to the behavioral room one day before the assays, where they were kept at 28 °C and were fed in the morning of the experiments. All behavioral assays were performed between 1–7 pm in a light and temperature-controlled cabinet (28 °C).

The swimming plus-maze (SPM) ^28^ consists of two shallow arms (length x width x height: 10 x 8 x 2.5 mm) and two deep arms (length x width x height: 10 x 8 x 5 mm) separated by a center zone. Six SPM arenas were glued onto an opaque acrylic platform illuminated from below by infrared LEDs. Fish were first pipetted from their home-tank to a well-plate and each of them was then placed in the center zone of a SPM arena. Fish activity was recorded for 5 minutes at 2 frames per second with a Basler ace2 camera placed 30 cm above the arena and equipped with a lens (FUJIAN; focal length=35 mm) covered with a long-pass infrared filter.

The movements of the animals were tracked and analyzed using Ethovision XT 17 ^51^. Mean velocity in the arena was extracted to assess locomotion. Time spent, frequency of entries, and latency to first entry in the shallow arms were extracted and z-scored. An anxiety score was then calculated by averaging z-scored latency, z-scored and sign-inverted time, and frequency measures to ensure consistent directionality to anxiety-like activity. Higher anxiety scores reflect increased anxiety-like behavior.

### Confocal microscopy

For single neurons’ projections and development experiments, fish were anesthetized in 0.1-0.2% tricaine (Merck, CAT: E10521-10G), embedded dorsal side-up in 2% low gelling point agarose (Merck, CAT: A9414-50g) and imaged under the confocal. For the imaging of broad DRN projections in cortisol and CUS experiments, 7 dpf Tg(*tph2:Gal4;UAS:Caax-GFP*) fish were euthanized and fixed overnight in a 4% PFA-0.25% Triton 100 X-100 solution at 4°C. Fish were rinsed and stored in PBS at 4°C until imaging.

A laser scanning confocal microscope (Zeiss LSM 700), equipped with water-dipping 10x (W N-Achroplan 10x/0.3 W M27) or 20x (W N-Achroplan 20x/0.5 W M27) objectives, was used for all recordings. Scans were recorded at 0.44-0.48 μm x 0.44-0.48 μm x 2.71 μm per pixel with the 20x objective, whereas images with the 10x objective were collected at 0.89 μm x 0.89 μm x 7.21 μm per pixel. For the 1 w.o. CUS experiments, close-up DRN scans were recorded at 0.69 μm x 0.69 μm x 2.71 μm. Laser wavelengths of 488 and 555 nm were used for fluorophore excitation. For experiments involving single-labelled neurons, laser intensity, digital gain, and offset were individually adjusted for each fish to optimize the signal-to-noise ratio. For the stress exposure experiments, these imaging parameters were kept constant across all samples to ensure valid comparisons between groups. Scans covered the brain dorso-ventrally and usually consisted of 110-180 slices. For images taken with the 20x objective, two scans were taken per fish to cover the whole brain from the olfactory bulb to the beginning of the spinal cord. The scans covering the anterior and posterior parts of each brain were stitched together using the Pairwise Stitching Plugin in FIJI ImageJ ^52^, using linear blending as fusion method and check peaks = 10.

### Reconstruction of projection patterns

#### Single neurons’ projections

Neurites of individual neurons were reconstructed semi-automatically using the Simple Neurite tracer (SNT) plugin in FIJI ImageJ ^53^. The neurites’ paths were then filled, binarized, and skeletonized in FIJI.

#### Broadly labelled DRN projections

Projections from broadly labelled DRN were too dense and numerous to be reconstructed using SNT. Using a single threshold over the whole scan to isolate the 5-HT fibers for each fish was not appropriate, since fluorescence intensity decreases along the dorso-ventral axis due to light scattering. To reconstruct the projectome in these broadly labelled fish, we first denoised the Tph2 channel in Fiji with a median filter (1-pixel radius). We then applied a custom MATLAB script to threshold the data, enabling user-defined selection of the threshold along the dorso-ventral axis. Threshold values were then applied over corresponding slices to generate a binary mask of labelled fibers. The experimenter was blind to scan identities until the completion of the analysis to avoid bias.

### Registration of Tph2 channels

Brain scans were registered to an age-matched common template brain using Computational Morphometry Toolkit (CMTK ^54^).

For registration of 1 w.o. scans, the command string (-T 16 -X 52 -C 8 -G 80 -R 3 -A ‘--accuracy 0.4’ -W ‘--accuracy 1.6’) was applied. Brain scans of Tg(*tph2:Gal4;UAS:Caax-GFP; gad1b:DsRed*) fish were registered to the Gad1:DsRed Z-brain template. Scans of Tg(*tph2:Gal4;UAS:Caax-GFP)* were saturated and registered to the Z-brain Reference Brain ^21^.

For registration of 2 w.o. scans, we first created a brain atlas at that development stage. A template brain was created by registering together several elavl3:H2b:GCaMP6s scans, obtained with the 10x objective at 0.59 µm x 0.59 µm x 7.64 µm per pixel, using the command string (-T 16 -X 52 -C 8 -G 80 -R 3 -A ‘--accuracy 0.4’ -W ‘--accuracy 1.6’) and averaging the four best registered scans. Template brains were then obtained for different transgenic lines (Tg*(gad1b:DsRed)*, Tg(*tph2:Gal4:UAS:Ntr-mCherry*), Tg(*pet1:mCherry*) and Tg(*tbx21:Gal4:UAS:tdTomato*)) after scanning the brains of fish outcrossed to the Tg(*elavl3:H2b:GCaMP6s*) line and registration to the elavl3H2b template using the same command string. We then registered the brain scans of the development, and chronic unpredictable stress experiments to the Gad1b:DsRed, and Tph2:Gal4:UAS:Ntr-mCherry template brains, respectively.

The accuracy of the registration procedure was carefully assessed for each fish by visually inspecting the overlap between the boundaries of the template brain and the registered brain. In addition, the alignment of several anatomical landmarks distributed along the dorsoventral axis was evaluated. For a subset of fish exhibiting suboptimal registration, a second registration attempt was performed after manually rotating the original brain scan to improve its alignment with the template prior to registration. This approach allowed several datasets to be successfully recovered. Fish that continued to show inaccurate registration after this procedure were excluded from further analyses.

### Determination of central nervous system regions innervated by Tph2 projections

For 1 w.o. fish, the Z-brain regions previously delineated by Randlett and colleagues ^21^ were used, with the two following exceptions. A shorter mask was created for the spinal cord to account for the differences in the posterior boundary of our scans compared to the Z-brain. A new mask was created for the medial zone of the dorsal telencephalon (Dm) based on anatomical landmarks of the elalv3 reference Z-brain. Amongst the 11 selected brain regions described throughout the results in this study, five of them (cerebellum, optic tectum-pretectum, hypothalamus, preoptic area, and thalamus) were built by agglomerating individual Z-brain regions together as described in Supp. table 6.

For 2 w.o. fish, the five main CNS subdivisions (telencephalon, mesencephalon, diencephalon, rhombencephalon, and spinal cord) and selected brain regions were manually delineated onto the 2 w.o. template brain using Fiji’s Segmentation Editor. Delineations were based on the distribution of the following markers (elavl3, gad1b, tph2, pet1, tbx21), as well as several resources describing the zebrafish brain across development ^55,56^.

The density of Tph2 projections to each region of interest was then obtained using custom MATLAB scripts by calculating the proportion of Tph2 labelled pixels within that region.

### Reconstruction and quantification of pruned and added neurites during the second week

The neurites pruned or added during the second week of development were identified by comparing the three-dimensional reconstructions of the same neuron at 1 w.o. and 2 w.o in SNT. The pruned neurites or new neurites’ paths were then filled, binarized, and skeletonized in FIJI.

To account for subtle variations in SNT semi-automatic path filling and ensure maximum overlap between the complete neurite tree and the corresponding reconstruction of pruned or new segments, all pruned and new traces were then dilated using FIJI’s 3D dilation plugin. The percentage of pruned and new neurites was calculated as follows for each fish:

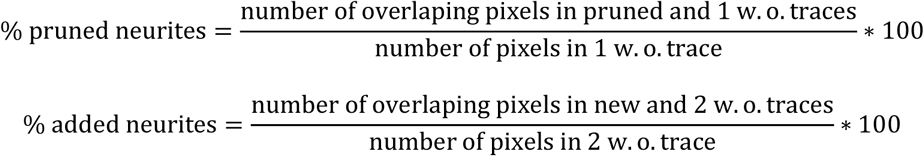

### Quantification of Tph2+ dorsal raphe nucleus size or Tph2+ DRN soma numbers

Due to overexposure of the DRN in the confocal stacks, resulting from the high laser power necessary to visualize ventral tph2:caax-GFP⁺ fibers, it was not possible to count individual DRN cell bodies in the 1 w.o. cortisol and 2 w.o. CUS experiments. Instead, a fixed intensity threshold was applied to all raw stacks’ GFP channel to generate binary masks of the Tph2+ DRN, and the area occupied by tph2⁺ cells was measured for each animal. Close-up, unsaturated DRN scans were taken for the 1 w.o. CUS experiment, which allowed counting Tph2+ DRN cells using the ROI manager tool in Fji.

### Statistical analysis

All statistical analysis were done in R statistical environment (R Core Team) or Graphpad Prism 10. Numerical data corresponding to graphs presented in the main and supplementary figures are provided in the Supplementary file ‘Numerical data.xls’. Statistical analyses pertaining to individual neurons projections were conducted with neurons treated as independent observations. A range of models (linear, generalized linear, and zero-inflation) was fitted to the data to describe the distribution of DRN fiber densities, and to categorize brain areas depending on the fiber density. Following BIC-value-based model selection, zero-inflation models with negative binomial distribution were applied. Area-projection categories were established based on the probability of zero DRN projection to a brain area (low or high), and on the level of non-zero density counts (low or high). Brain areas with low zero-density probability and high level of non-zero count are considered densely innervated by many neurons, while areas with high zero-density probability and low level non-zero count are considered densely innervated by few neurons. Brain areas with high zero probability and high non-zero count are considered sparsely innervated areas by few neurons.

Spearman correlations were used to determine the relationship between soma location, or Gad1b phenotype, and the organization of neurons’ projections along the antero-posterior and dorso-ventral axes. Multiple Spearman correlation analyses were performed on all Tph2^+^ neurons, or separately on Tph2^+^GAD1b^-^ and Tph2^+^GAD1b^+^ neurons, to assess the relationship between the soma location and the density of fibers within brain areas. P-values were adjusted using the Benjamini-Hochberg method. For the longitudinal study, pairwise t-tests were conducted to assess the change in projection and laterality scores from the first to the second week of development, as well as to compare the overall percentages of neurite loss and neurite growth. To investigate whether the projections undergo more changes in certain CNS regions, mixed-effects models (REML) were fitted, followed by Benjamini-Hochberg FDR correction in case of a significant effect.

For the analysis of SPM assays in the stress exposure experiments, a two-level outlier discrimination was done. Subjects were excluded if they exhibited extreme values in both locomotion and anxiety score, defined as values below the lower quartile minus the interquartile range or above the upper quartile plus the interquartile range. Following exclusion, anxiety-like variables were rescaled. To assess the relationship between behavioral variables and hydrocortisone-induced stress, linear models were fitted. Pairwise post-hoc comparisons of control and each treated group were done using Dunnet’s correction. To assess the relationship between behavioral variables and chronic unpredictable stress, two-sided student t-tests were conducted. Pairwise comparisons of anxiety scores were conducted using one-sided tests in both experiments, based on the a priori hypothesis that stressor exposure increases anxiety ^27,29,45^.

To assess the effect of either hydrocortisone-induced stress during the first week or chronic unpredictable stress during the second week on fiber density in different brain areas, two-way repeated measures ANOVA (tw-rANOVA) models were fitted, with the brain areas and the stress exposure as fixed factors and the subject identifier as repeated factors. Regions containing tph2+ cell bodies (pons-medulla and spinal cord) were excluded from the analysis to ensure that the measured signal reflected neurite projections rather than somatic labeling. In case of a significant interaction between the area identifier and the stress exposure, pairwise post-hoc comparisons were done using Benjamini-Hochberg FDR correction. To test the hypothesis that chronic unpredictable stress treatment during the first week would increase fiber density in the same telencephalic areas as with the cortisol treatment, one-tailed student t-tests were performed between control and stressed groups in the olfactory bulbs, subpallium, pallium, and Dm. To determine the effect of stress exposure on the number of tph2+ DRN cells, the total tph2+ pixel count within the DRN was compared between control and treatment groups using a one-way ANOVA for the hydrocortisone experiment and a two-sided student t-test for the 2 w.o. chronic unpredictable stress experiment. For the 1 w.o. chronic unpredictable stress experiment, a two-sided student t-test was conducted to assess whether the number of Tph2+ DRN neurons differed across groups.

## AUTHORS CONTRIBUTIONS

L.J.F. and F.K conceived the study; F.K. supervised and acquired funding. L.J.F. conducted most experiments, with help from Z.K.V, K.T. and F.K.; L.J.F., Z.K.V, K.T. and F.K. analyzed the data. L.J.F prepared the figures; L.J.F. and F.K. wrote the original draft; L.J.F., Z.K.V., K.T. and F.K. reviewed and edited the manuscript.

## ACKNOWLEDGMENTS

We thank colleagues for generously sharing transgenic lines: Dr. Harold Burgess (Tg(Tph2:Gal4:UAS-Ntr-mCherry) line), Dr. Aristide Arrenberg (Tg(UAS:Caax-GFP) line), Dr. Shin-ichi Higashijima (Tg(gad1b: DsRed) line), Dr. Christiana Lillesaar (Tg(Pet1a:mCherry) line). We thank the expert staff from the Core Facility for Integrated Bioimaging (CFIM/CFIB) at UCPH for training and guidance with microscopy image acquisition and analysis. This work was funded by the Lundbeck Foundation (Ascending Investigator grant to F.K.).

